# Defective splicing of Y-chromosome-linked gigantic genes contributes to hybrid male sterility in Drosophila

**DOI:** 10.1101/2025.05.22.655455

**Authors:** Adrienne Fontan, Romain Lannes, Jaclyn M. Fingerhut, Jullien M. Flynn, Yukiko M. Yamashita

**Affiliations:** Whitehead Institute for Biomedical Research; Massachusetts Institute of Technology, Department of Biology; Howard Hughes Medical Institute

## Abstract

The Y chromosome evolves rapidly, often differing dramatically even between closely related species. While such divergence has long been suspected to contribute to hybrid male sterility, leading to reproductive isolation and thus speciation, the underlying mechanisms remain elusive. Here, we identify a molecular basis linking Y chromosome divergence to reproductive isolation in *Drosophila*. We show that male hybrids between *D. simulans* and *D. mauritiana* fail to properly express key Y-linked fertility genes. These genes contain unusually large introns, exceeding megabases and show substantial sequence divergence between species. In the hybrids, these gigantic introns are misprocessed, resulting in widespread splicing defects, including aberrant “back-splicing” events that join later exons to earlier ones. Our findings suggest that sequence divergence within introns can disrupt essential gene expression through defective splicing, providing a mechanistic link between rapid Y chromosome evolution and hybrid sterility. This work highlights the underappreciated role of intronic divergence in speciation.

## Introduction

Extensive studies have provided insights into the process of reproductive isolation, which prevents gene flow between two populations, leading to speciation (Peichel, et al. 2024). Although there are multiple causes of reproductive isolation, one that often appears early in the course of speciation is hybrid sterility (Presgraves and Meiklejohn 2021). The heterogametic sex (XY males and ZW females) suffers hybrid sterility and inviability more often than the homogametic sex (XX females and ZZ males), an observation known as Haldane’s rule (Haldane 1922). Multiple hypotheses have been put forward to explain Haldane’s rule (Orr 1997; Delph and Demuth 2016). For example, the dominance theory proposes that the heterogametic sex, which has only one X chromosome from one species, may suffer recessive incompatibility with the autosomes from the other species, leading to inviability or sterility in hybrids (Turelli and Orr 1995). It has also been hypothesized that incompatible interactions between two sex chromosomes (X vs. Y or Z vs. W) from different species may explain Haldane’s rule (Delph and Demuth 2016). It is generally thought that Haldane’s rule likely represents a composite phenomenon resulting from multiple causes (Orr 1997; Cutter 2024). Accordingly, much remains to be understood about how individual factors contribute to hybrid sterility in a manner that preferentially impacts the heterogametic sex.

While many studies have documented hybrid sterility across species including mammals, insects, and plants, Drosophila species have served as a leading model of hybrid sterility for a century (Presgraves 2008; Coughlan and Matute 2020; Searle and Pardo-Manuel de Villena 2024), including the three *D. simulans* clade species (*D. simulans, D. mauritiana*, and *D. sechellia)* that diverged ∼250 thousand years ago (Fig 1A) (Lachaise, et al. 1986; Johnson, et al. 1993; Presgraves and Meiklejohn 2021). These species exhibit hybrid male sterility in reciprocal crosses (Lachaise, et al. 1986), whereas hybrid females are fertile, conforming to Haldane’s rule (Haldane 1922). Mapping of the genes that cause sterility of *D. simulans/D. mauritiana* hybrid males has revealed the presence of multiple factors on the X chromosome and the autosomes (Coyne and Charlesworth 1986; Cabot, et al. 1994; Palopoli and Wu 1994; Hollocher and Wu 1996; True, et al. 1996). One such genes, the X-linked *OdsH* gene, is sufficient to cause male sterility in a heterospecific context: *D. simulans* males with the *D. mauritiana OdsH* gene are sterile (Perez, et al. 1993; Perez and Wu 1995). Cytologically, *D. simulans/D. mauritiana* hybrid males exhibit abnormalities during spermatid differentiation, such as defects in axoneme elongation and sperm individualization (Lachaise, et al. 1986; Kulathinal and Singh 1998; Kanippayoor, et al. 2020). Several autosomal genes required for spermatogenesis (e.g., *sa, dj, Mst84Dc,* and *Mst98Ca*) were downregulated in *D. simulans/D. mauritiana* hybrids (Michalak and Noor 2003, 2004; Moehring, et al. 2007; Catron and Noor 2008; Sundararajan and Civetta 2011), potentially explaining the cytological phenotype observed in the sterile hybrids.

**Figure 1:**
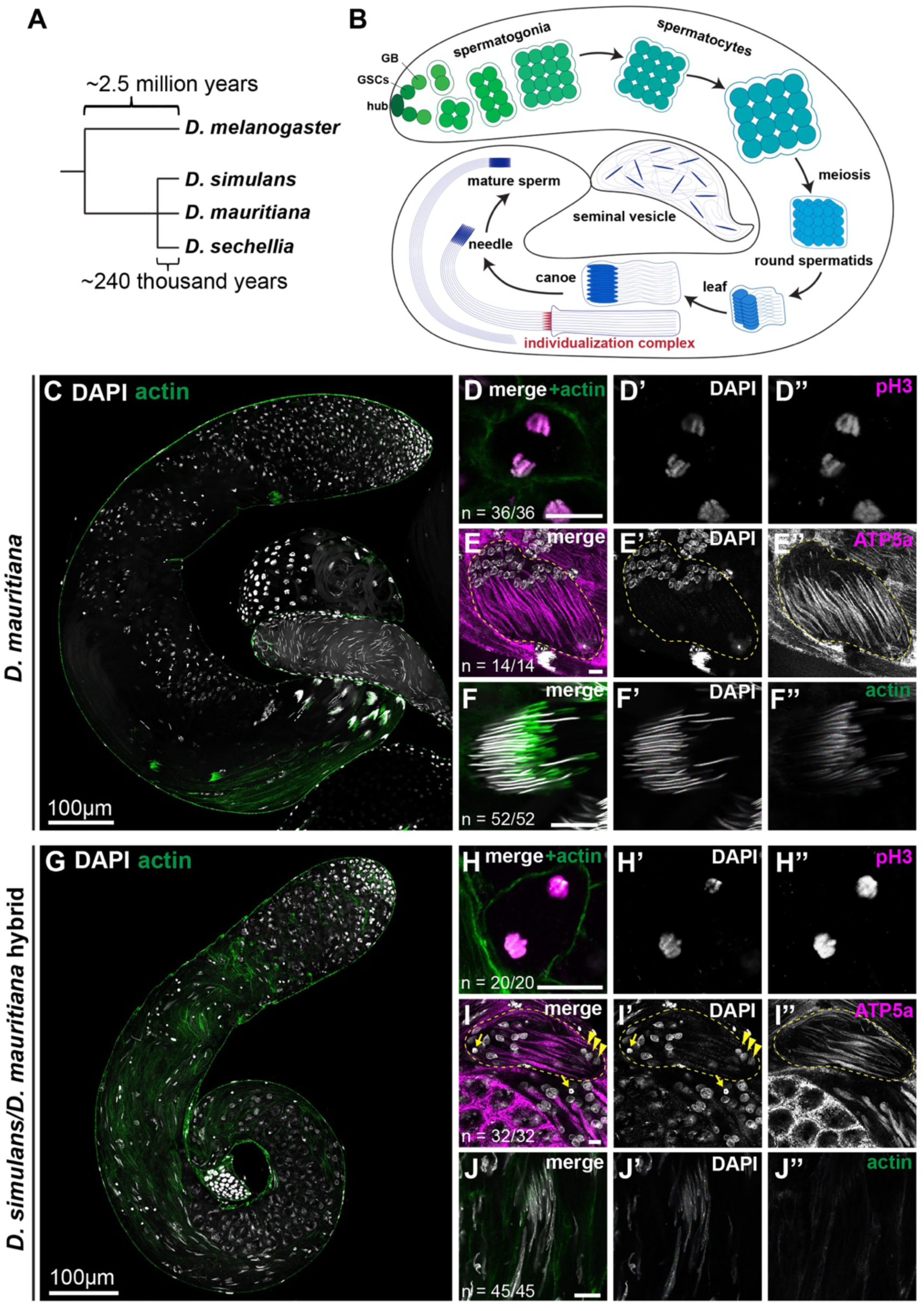
*D. simulans/D. mauritiana* hybrids display cytological defects in post-meiotic spermatids. A. Phylogeny of the *Drosophila melanogaster* complex. B. Schematic of *Drosophila* spermatogenesis. Germline stem cells reside in the niche (formed by hub cells) at the apical tip of the testis. Differentiating progeny of germline stem cells undergo transit-amplifying divisions as gonialblasts (GB) and then as spermatogonia with incomplete cytokinesis, yielding a cyst of 16 interconnected germ cells. These cells enter the spermatocyte stage, growing in size and expressing genes required for meiosis and post-meiotic development. Spermatocytes then undergo meiosis, and haploid spermatids undergo morphological transformation to become functional sperm. C-F. *D. mauritiana* testis and germ cells. Low magnification view of the whole testis (C) stained for F-actin (green, stained with phalloidin-Alexa 488) and DAPI (gray). Meiotic anaphase (D) stained for phospho-H3 ser10 (magenta) and DAPI (gray). Early elongating spermatids (E) stained for ATP5a (magenta, mitochondria) and DAPI (gray). Needle stage spermatids (F) stained for F-actin (green, stained with phalloidin-Alexa 488, individualization complex). Scale bar: 10µm, unless otherwise noted. G-J. *D. simulans/D. mauritiana* hybrid testis and germ cells. Low magnification view of the whole testis (G), meiotic anaphase (H), early elongating spermatids (I), and canoe stage spermatids (J), with the same staining as C-F. Scale bar: 10µm, unless otherwise noted. Arrows indicate hyper-condensed (dying) nuclei; arrowheads indicate depolarized spermatid nuclei.

In parallel, the Y chromosome, essential for male fertility in Drosophila (Bridges 1916), has been speculated to contribute to hybrid male sterility. The Drosophila Y chromosome has diverged rapidly, as is the case for many species, including mammals (Hughes and Page 2015; Kotov, et al. 2022), leading to a model that Y chromosome divergence may contribute to the hybrid male sterility observed in Drosophila species. Indeed, introgression of the Y chromosome into other species (e.g., the *D. simulans* Y chromosome in a *D. mauritiana* background and vice versa) causes sterility, suggesting that the Y chromosome is not compatible between these species (Johnson, et al. 1993; Zeng and Singh 1993; Araripe, et al. 2016). However, the underlying molecular mechanisms that render the Y chromosome incompatible in hybrids remain poorly understood.

In recent years, Y chromosome sequence divergence between Drosophila species has been characterized in more detail, revealing the rearrangement of gene locations on the Y chromosomes and considerable changes in repetitive DNA sequences, such as satellite DNA (Jagannathan, et al. 2017; Chang, et al. 2022). However, functional divergence between Y chromosomes is poorly understood. Satellite DNA divergence has been speculated to underlie hybrid incompatibility, although the molecular mechanisms behind it are still not fully understood (Yunis and Yasmineh 1971; Presgraves and Meiklejohn 2021). Curiously, satellite DNA is found in the introns of several Y-linked fertility genes. These genes have satellite DNA-containing gigantic introns sometimes exceeding megabases in size, while protein-coding sequences (exons) are only up to ∼15kb in total, a phenomenon called intron gigantism (Kurek, et al. 1996; Carvalho, et al. 2000; Kurek, et al. 2000). Intron gigantism is observed in a handful of genes within the genomes of a broad range of species, from flies to humans: a well-known example is the mammalian dystrophin gene, which spans over 2 megabases but encodes only 11kb of the protein-coding sequence (Pozzoli, et al. 2002; Pozzoli, et al. 2003). While the functional significance of intron gigantism remains unclear, studies of Y-linked fertility genes in *D. melanogaster* suggest that these enormous introns present substantial obstacles to gene expression, particularly during transcription and splicing (Fingerhut, et al. 2019; Fingerhut, et al. 2024). For example, the transcription of gigantic genes attenuated within gigantic introns upon perturbation of splicing (Fingerhut, et al. 2024).

Here, we show that hybrid males produced by the mating of *D. simulans* females and *D. mauritiana* males exhibit unique defects in the expression of Y-linked gigantic genes, likely contributing to hybrid sterility. Our analyses reveal that the compromised expression of fertility genes in hybrids results, at least in part, from splicing errors particularly at gigantic introns. DNA sequences of gigantic introns are poorly conserved between *D. simulans* and *D. mauritiana*. Taken together, we propose that the defective splicing of gigantic introns due to poorly conserved intronic sequences contributes to hybrid male sterility. Our results may provide a mechanistic link between the rapid divergence of the Y chromosome and hybrid male sterility, providing a potential explanation for Haldane’s rule.

## Results

### Defective spermatogenesis of *D. simulans/D. mauritiana* hybrids is associated with downregulation of Y-chromosome-linked fertility genes

Interspecies crosses between *D. simulans* females and *D. mauritiana* males have long been known to produce sterile hybrid males due to defects in post-meiotic processes, including sperm axoneme development and sperm DNA packaging (Lachaise, et al. 1986; Kulathinal and Singh 1998; Kanippayoor, et al. 2020) (Fig 1). Spermatogenesis of the parental species, *D. simulans* and *D. mauritiana,* is highly similar in gross anatomy and cellular features to the well-characterized *D. melanogaster* spermatogenesis, as described previously (Fuller 1993; Yamashita 2018) (Fig 1B-F, Supplementary Fig S1). The apical tip of the testis contains mitotically proliferating cells (germline stem cells, gonialblasts, and spermatogonia), which undergo four rounds of mitosis with incomplete cytokinesis to yield a cluster of 16 spermatogonia. These cells then enter the spermatocyte stage, progressively growing in size (Fig 1C, Supplementary Fig S1A). Mature spermatocytes undergo meiosis (Fig 1D, Supplementary Fig S1B), and the resulting 64 interconnected haploid spermatids morphologically transform in a cyst (Fig 1E, Supplementary Fig S1C): the nuclei undergo stereotypical morphological changes through leaf, canoe, and needle stages, resulting in highly condensed sperm DNA (Tokuyasu 1974; Fabian and Brill 2012)(Fig 1E, F, Supplementary Fig S1C, D). The entire cellular morphology also undergoes striking morphological changes, elongating the sperm axoneme and surrounding mitochondria to form 1.9mm sperm tails (Fig 1B). An actin-based structure called the individualization complex forms around needle-stage spermatid nuclei (Fig 1F, Supplementary Fig S1D), which subsequently progresses downwards toward sperm tails to eliminate excess cytoplasm and produce individual sperm (Tokuyasu, et al. 1972; Fabian and Brill 2012). Mature sperm are then released into the seminal vesicle.

In contrast to the parental species, *D. simulans/D. mauritiana* hybrid males exhibited various defects in post-meiotic stages, as described previously (Lachaise, et al. 1986; Kulathinal and Singh 1998; Kanippayoor, et al. 2020). Although they possessed normal gross anatomy with the stereotypical coiled tubular structure (Fig 1G), and underwent apparently normal germ cell development until meiotic divisions (Fig 1H), the hybrids began to exhibit clear defects after meiosis. First, in contrast to the parental species, where all 64 nuclei within a cyst polarize uniformly towards the distal end of the testis (Fig 1E, Supplementary Fig S1C), the hybrid cyst displayed depolarization with some nuclei incorrectly polarized towards the opposite end (Fig 1I, arrowheads). Some nuclei were hyper-condensed, indicating cell death (Fig 1I, arrows). Spermatids ultimately failed to condense their nuclear DNA properly (Fig 1J), never forming the needle-shaped nuclei of mature sperm observed in parental species (Fig 1F, Supplementary Fig S1D). Furthermore, the individualization complex (Fig 1F, Supplementary Fig S1D) did not form in hybrids (Fig 1J). These cytological defects (described previously and further detailed in this study) explain the well-known sterility of *D. simulans/D. mauritiana* hybrids.

RNA sequencing on the testes of hybrids compared to those of the parental species revealed the downregulation of several Y-linked genes (Table 1). We conducted RNA sequencing using total RNA to capture transcription intermediates in addition to mature mRNAs. The results confirmed the previously reported downregulation of several autosomal genes in hybrids (see Methods). We then focused on Y-linked genes, because the Y chromosome harbors genes required for post-meiotic stages of spermatogenesis (Gatti and Pimpinelli 1992), and it has been shown that introgression of heterospecific Y chromosomes into other species’ backgrounds led to sterility (Johnson, et al. 1993; Zeng and Singh 1993). Because the hybrid males studied here carry the *D. mauritiana* Y chromosome, the hybrids’ RNA sequence reads were aligned to the *D. mauritiana* Y chromosome assembly (Chang, et al. 2022) and were compared to the transcriptome of *D. mauritiana* testes (Supplementary_file_1, Methods). Differential expression analysis showed that a subset of Y-linked genes, most notably *kl-3* and *kl-5*, were downregulated in hybrids (Table 1). We observed a slight but significant reduction in the expression of *kl-2*, *WDY*, *ppr-Y*, and *CCY* as well.

**Table 1:** Differential gene expression analysis of Y-linked genes in *D. simulans/D. mauritiana* hybrids. *D. simulans/D. mauritiana* hybrids vs. *D. mauritiana* differential Y genes expression results. Blue indicates statistically overexpressed genes in the *D. simulans/D. mauritiana* hybrids. Red indicates genes statistically under expressed in the *D. simulans/D. mauritiana* hybrids. Darker red highlight genes statistically under expressed in the *D. simulans/D. mauritiana* hybrids with log2 fold change < −1.

| FBgn ID | baseMean | log2FoldChange | pvalue | padj | symbol |
| --- | --- | --- | --- | --- | --- |
| FBgn0267433 | 16779.8014 | -1.317262 | 8.88E-45 | 6.46E-43 | kl-5 |
| FBgn0267432 | 15681.8204 | -1.208128 | 1.42E-47 | 1.15E-45 | kl-3 |
| FBgn0001313 | 7835.85713 | -0.760683 | 5.04E-15 | 6.88E-14 | kl-2 |
| FBgn0267449 | 2896.11389 | -0.715952 | 4.43E-09 | 3.14E-08 | WDY |
| FBgn0046697 | 700.079317 | -0.697227 | 8.42E-06 | 3.81E-05 | Ppr-Y |
| FBgn0261399 | 65.240147 | -0.687174 | 4.22E-02 | 8.29E-02 | Pp1-Y1 |
| FBgn0267592 | 5574.79779 | -0.278345 | 1.51E-02 | 3.37E-02 | CCY |
| FBgn0265047 | 11838.8318 | -0.147742 | 9.82E-02 | 1.69E-01 | FDY |
| FBgn0046323 | 4764.88505 | -0.024502 | 7.87E-01 | 8.51E-01 | ORY |
| FBgn0267490 | 117772.255 | 0.010969 | 8.97E-01 | 9.29E-01 | MST77Y-4 |
| FBgn0058064 | 2819.95806 | 0.36048 | 2.95E-04 | 1.00E-03 | ARY |
| FBgn0046698 | 849.1199 | 0.718349 | 2.92E-06 | 1.41E-05 | Pp1-Y2 |
| FBgn0267489_alt | 1234.04319 | 0.829099 | 5.78E-13 | 6.29E-12 | PRY_alt |

The *kl-2*, *kl-3*, and *kl-5* genes encode axonemal dynein subunits, which are required for sperm tail development and thus fertility (Carvalho, et al. 2000). Using single molecule RNA FISH (smRNA FISH) with probes for *kl-3* exons 1 and 14, we monitored the expression of *kl-3* in *D. simulans/D. mauritiana* hybrids over the course of spermatocyte development. Because the *kl-2*, *kl-3*, and *kl-5* genes contain gigantic introns that exceed megabases in length, their expression takes the entirety of spermatocyte development (80-90 hours) (Chandley and Bateman 1962; Fingerhut, et al. 2019), where the transcription of early exons begins in early spermatocytes, followed by the transcription of gigantic introns, then concludes with transcription of later exons in mature spermatocytes (Fingerhut, et al. 2019). The mature mRNAs of *kl-2, kl-3* and *kl-5* are observed as cytoplasmic RNP granules, termed kl-granules, which contain Pontin protein (Fingerhut, et al. 2019; Fingerhut and Yamashita 2020). Expression of *kl-3* in *D. simulans* and *D. mauritiana* exhibited a spatiotemporal pattern similar to that observed in *D. melanogaster* (Fig 2B, D) (Fingerhut, et al. 2019): the transcription of exon 1 was observed in early spermatocytes, whereas exon 14 was transcribed much later, concluding with the formation of kl-granules in the cytoplasm (Fig 2C, E), which also contained a known protein component, Pontin (Fig 2H, I)(Fingerhut and Yamashita 2020).

**Figure 2:**
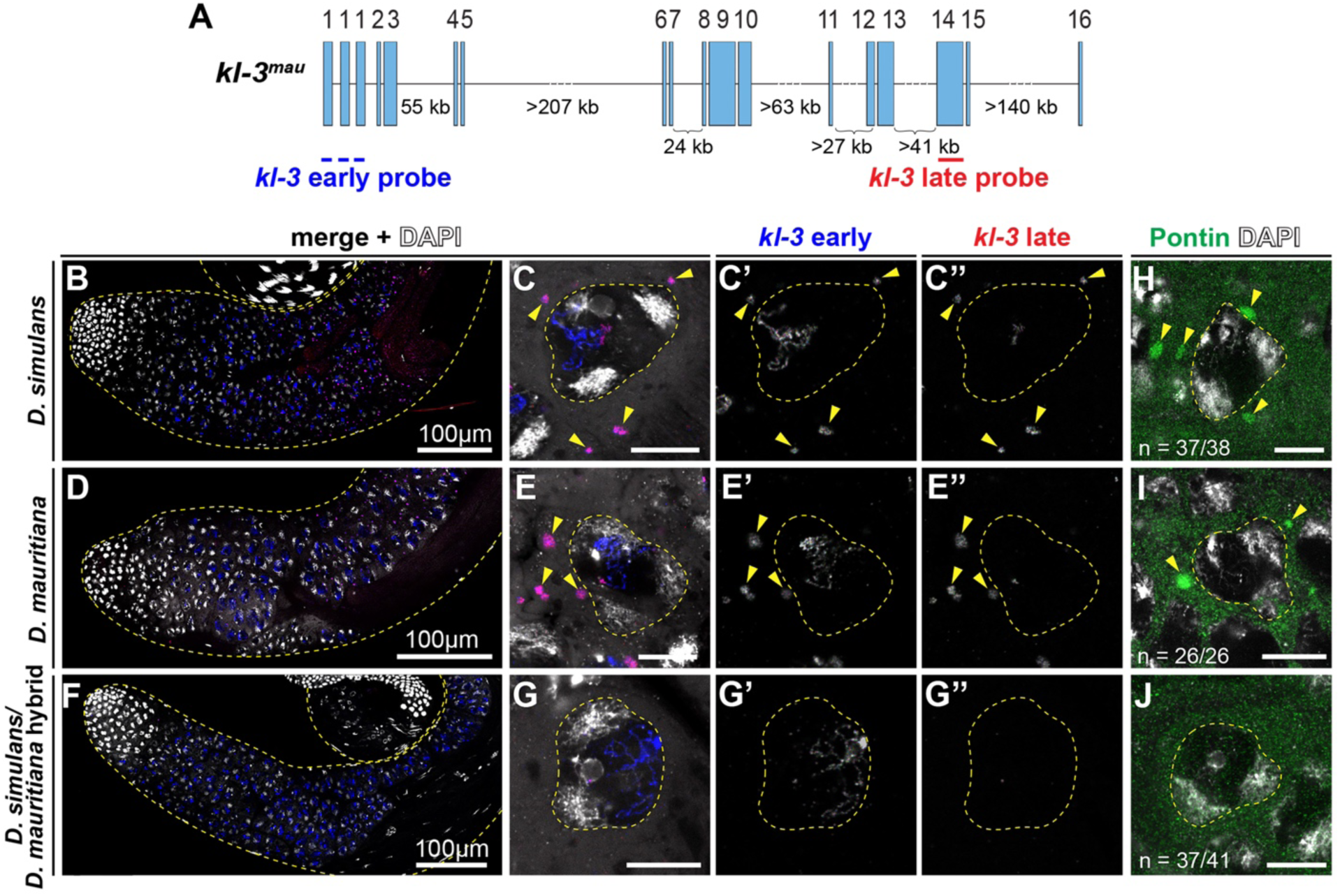
kl-granules are absent from *D. simulans/D. mauritiana* hybrid spermatocytes. A. *D. mauritiana kl-3* (*kl-3^mau^*) gene structure. The positions of smRNA FISH probes are indicated. B-G. RNA in situ hybridization for *kl-3* transcripts in *D. simulans* (B, C), *D. mauritiana* (D, E) and *D. simulans/D. mauritiana* hybrids (F, G). Apical third of the testis (B, D, F) and individual late spermatocyte nuclei (C, E, G) are shown. Spermatocyte nuclei are indicated by dashed lines. Arrowheads indicate cytoplasmic kl-granule containing mature *kl-3* mRNA. Blue: *kl-3* early exons. Red: *kl-3* late exons. Gray: DAPI. Scale bar: 10µm, unless otherwise noted. H-J. Individual late spermatocyte nucleus from *D. simulans* (H), *D. mauritiana* (I) and *D. simulans/D. mauritiana* hybrids (J). Spermatocyte nuclei are indicated by dashed lines. Arrowheads indicate cytoplasmic kl-granule stained for anti-Pontin antibody (green).

In contrast to *D. simulans* and *D. mauritiana*, we found that *D. simulans/D. mauritiana* hybrids exhibit striking defects in *kl-3* expression. Although exon 1 transcripts appeared in the early spermatocyte stage, exon 14 was rarely visible, and cytoplasmic kl-granules were absent, suggesting functional *kl-3* mRNA was not produced (Fig 2F, G). Similar results were obtained for the *kl-2* and *kl-5* genes, where the parental species exhibited the same expression patterns as *D. melanogaster* (Supplementary Fig S2B, C, E, F), but the hybrids did not (Supplementary Fig S2D, G). As expected from the lack of cytoplasmic *kl-2*, *kl-3*, and *kl-5* transcripts, immunofluorescence staining for Pontin showed that kl-granules were indeed absent in the hybrids (Fig 2J). These results demonstrate that hybrids fail to produce mature mRNA of *kl-2, kl-3* and *kl-5* transcripts. We conclude that *D. simulans/D. mauritiana* hybrid males fail to express a subset of Y-linked axonemal dynein genes, which may contribute to hybrid sterility.

### Transcription of *kl-2*, *kl-3,* and *kl-5* attenuates within gigantic introns in hybrids

More detailed analysis of the RNA sequencing reads revealed that the expression of *kl-2*, *kl-3*, and *kl-5* was specifically decreased toward the 3’ side of the gene. For example, the read depth of *kl-3* dropped dramatically between exons 5 and 6 in hybrids relative *to D. mauritiana* (Fig 3A). Notably, the intron between exons 5 and 6 is a gigantic intron, containing 207kb of assembled sequence and a gap in the genome assembly due to the presence of repetitive DNA (Supplementary_file_1)(Chang, et al. 2022). A similar drop was observed between exons 13 and 14 of *kl-5,* and between exons 6 and 7 of *kl-2*, which are both gigantic introns (35kb plus a gap and 432kb plus a gap, respectively) (Fig 3A, Supplementary_file_1). These results are consistent with the absence of late exon of *kl-2*, *kl-3*, and *kl-5* visualized by smRNA FISH (Fig 2F, G, Supplementary Fig S2D, G). Together, these results revealed that *kl-2*, *kl-3*, and *kl-5* become downregulated in hybrid testes due to transcription attenuation within gigantic introns.

**Figure 3:**
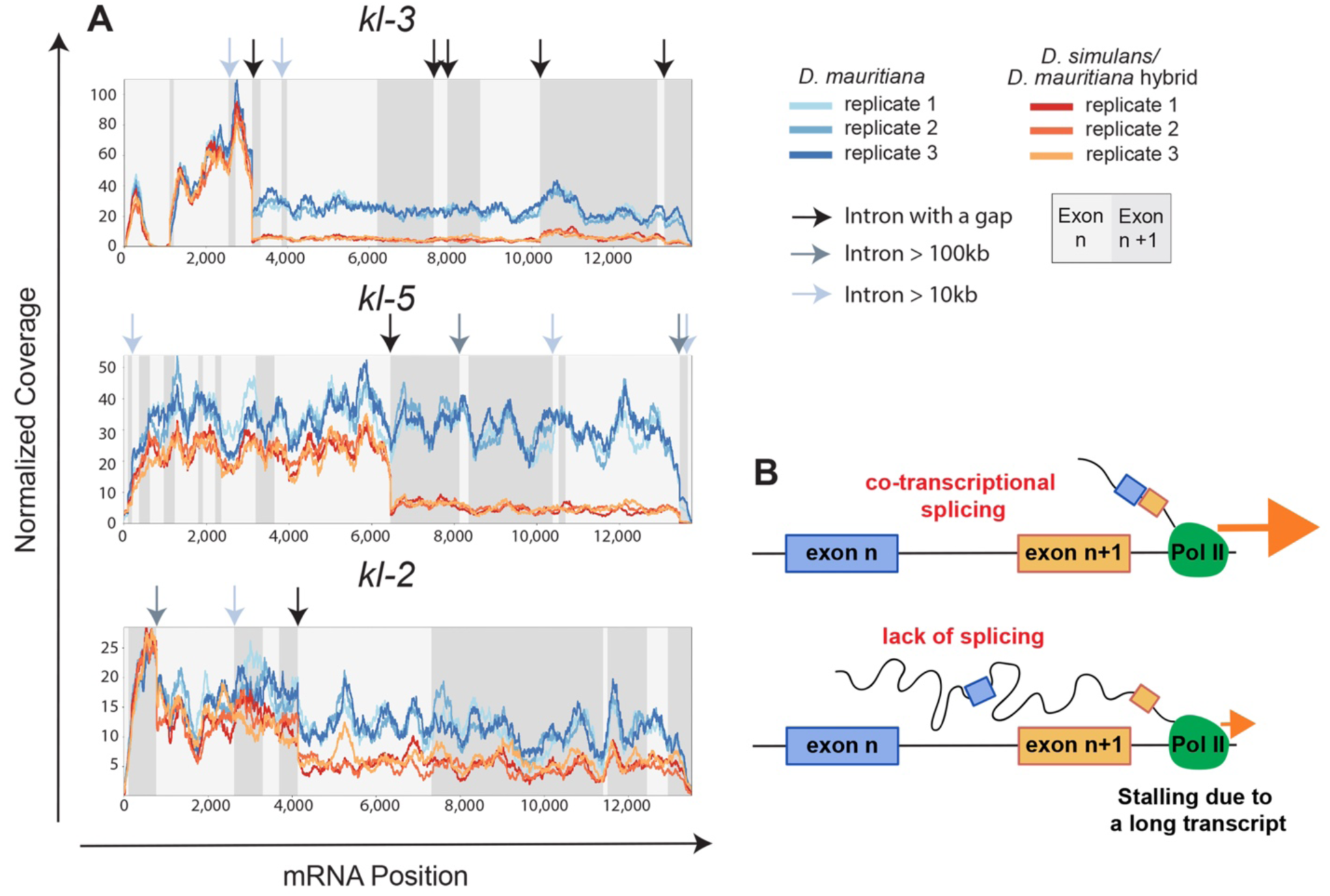
Transcription of *kl-2, kl-3,* and *kl-5* attenuates in *D. simulans/D. mauritiana* hybrids relative to *D. mauritiana* within gap-containing gigantic introns. A. Normalized read depth coverage of the *kl-2, kl-3,* and *kl-5* genes along the coding sequence in *D. mauritiana* vs. *D. simulans/D. mauritiana* hybrids. Alternating shades of gray in the background indicate different exons. Arrows point to the location of large or gap-containing introns. B. A proposed model of stalled transcription due to defective splicing (after (Fingerhut, et al. 2024)).

The sudden drop in read depth within gigantic introns is reminiscent of a phenotype we recently discovered in *D. melanogaster* following depletion of splicing factors (Fingerhut, et al. 2024). We have shown that these gigantic introns are co-transcriptionally spliced, and co-transcriptional splicing is critical for the continuation of transcription (Fingerhut, et al. 2024). We postulated that transcription may attenuate within gigantic introns upon perturbation of splicing, perhaps because excessively long unspliced transcripts attached to RNA polymerase II hinder the progression of transcription (Fig 3B, see also an alternative interpretation below) (Fingerhut, et al. 2024). The dependence of transcription on splicing was specific to genes containing gigantic introns (Fingerhut, et al. 2024), suggesting large introns are more sensitive to splicing perturbation. Considering these similarities, we hypothesized that Y-linked genes with gigantic introns may be downregulated in hybrids due to splicing defects.

### *D. simulans/D. mauritiana* hybrids are defective in the splicing of gigantic introns

To test whether the read depth drop observed in the *D. simulans/D. mauritiana* hybrids may be associated with splicing defects, we analyzed the splicing of *kl-2*, *kl-3*, and *kl-5* in *D. simulans/D. mauritiana* hybrids compared to *D. mauritiana*. To this end, we developed a bioinformatics pipeline that examines not only the efficiency of splicing (i.e., the number of correctly spliced reads divided by the total number of spliced and nascent reads) but also potentially incorrect splicing events in which the end of an exon is connected to an unpredicted sequence (see Methods). Using this method, all reads containing the 3’ end of a given exon were identified, and the identity of the adjoining sequence was analyzed. These reads were categorized as follows (Fig 4A):

i. Spliced: the 3’ end of the exon was correctly joined to the 5’ end of the next exon.
ii. Unspliced: the junction between the 3’ end of the exon and the juxtaposing intron remained intact, indicative of a nascent RNA.
iii. Exon-other: the 3’ end of the exon was connected to a downstream sequence that was not the predicted splice acceptor site. These reads may represent erroneous splicing events or products of recursive splicing. (Note that the reads that correspond to known alternative splice isoforms or overlapping gene were excluded from this category).
iv. Exon-clipped: the read was partially mapped to the 3’ end of the exon, but the adjoining sequence could not be mapped linearly with the genome, and was marked as clipped by the aligner. This category of reads is indicative of chimeric fragments between 3’ end of an exon and an RNA fragment that is not normally expected to be joined to the given exon. Such chimera may be biological, or also arise from issues with the library quality, such as the artifactual ligation of two unrelated DNA molecules during library preparation. Therefore, these reads require additional analysis for validation. Note that RNA sequence reads were considered in this category only if the ‘clipped’ portion was greater than 10 nucleotides.

**Figure 4.**
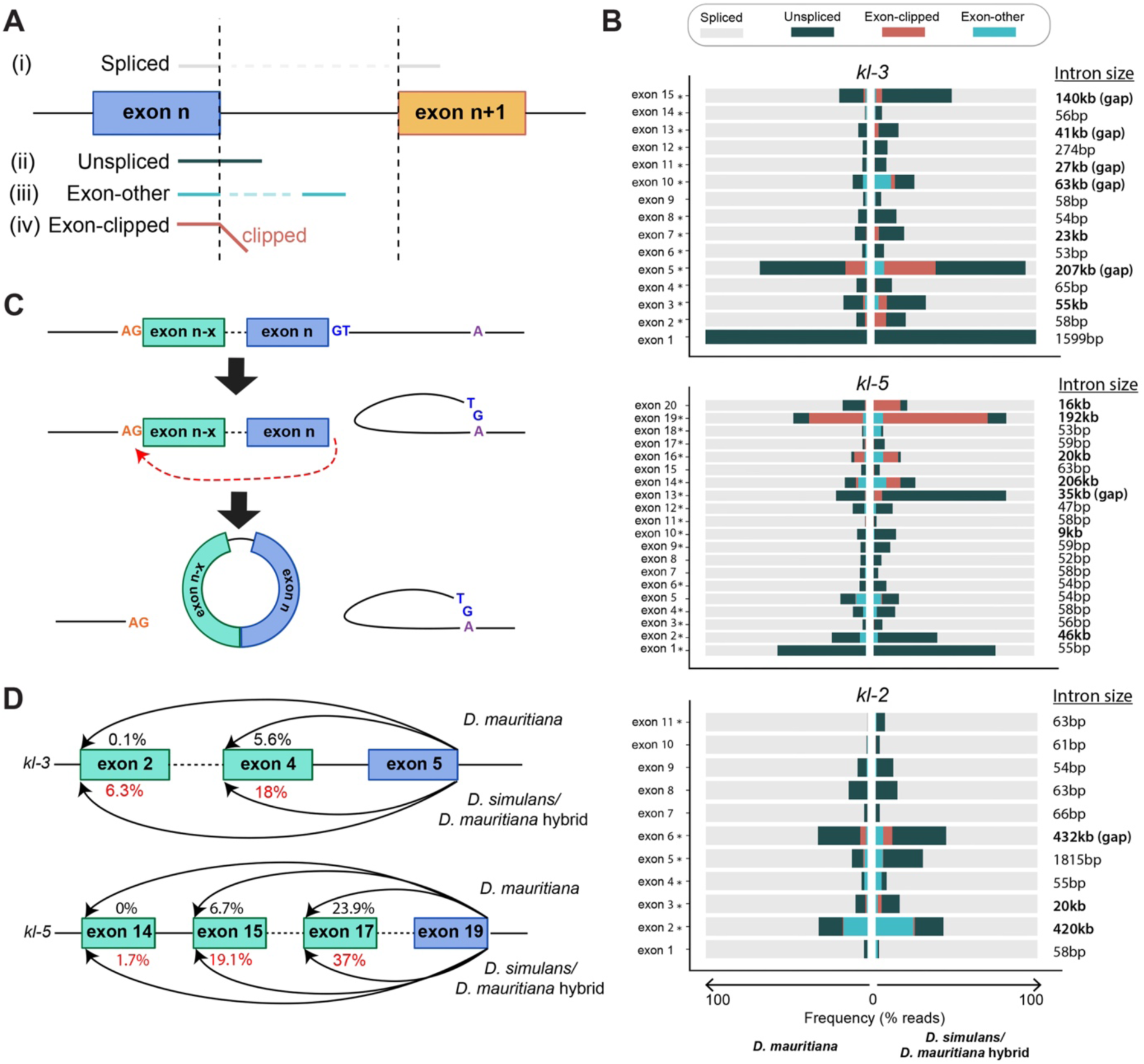
*D. simulans/D. mauritiana* hybrids exhibit splicing defects in *kl-2, kl-3,* and *kl-5*. A. Categories of sequence reads that overlap with the end of exons. B. Sequence read analysis for each exon end of the *kl-2, kl-3,* and *kl-5* genes in *D. mauritiana* vs. *D. simulans/D. mauritiana* hybrids. Asterisks(*) indicate significant difference in the proportion of correctly spliced reads between *D. mauritiana* and *D. simulans/D. mauritiana* hybrids. C. Schematic of back-splicing producing circular RNA. D. Frequency of back-spliced reads for *kl-3* exon 5 ends and *kl-5* exon 19 ends in *D. mauritiana* vs. *D. simulans/D. mauritiana* hybrids. Percentages indicate the frequency within the total reads aligning to the end of *kl-3* exon 5 and *kl-5* exon 19.

We performed this analysis for each exon of *kl-2, kl-3,* and *kl-5* (Fig 4B). Several locations displayed significant difference between *D. mauritiana* and *D. simulans/D. mauritiana* hybrids (Fig 4B, introns marked with asterisks, see also Supplementary file_2, and Supplementary file_3), including *kl-3* exons 5 and 15, and *kl-5* exons 13 and 19, all of which immediately precede gigantic introns. The sequence reads across these junctions exhibited several notable features. First, the proportion of correctly spliced products decreased considerably in the hybrids, compared to *D. mauritiana*. Correspondingly, the proportion of unspliced reads often increased in the hybrids, suggesting that hybrids indeed fail to splice gigantic introns. It should be noted that even parental species (*D. mauritiana*) often showed a large fraction of unspliced reads, likely reflecting the early spermatocyte population that had not transcribed the next exon or splice acceptor site.

Most notably, we found a considerable increase in exon-clipped reads in hybrids, where the 3’ end of an exon (particularly, *kl-3* exon 5, *kl-5* exon 14, and *kl-5* exon 19, all of which immediately precede gigantic introns) was joined to ‘unexpected’ sequence (Fig 4A, B). Because exon-clipped reads are normally assumed to be a result of experimental artifacts, they are frequently excluded from analyses. However, given the possibility of splicing defects in hybrids, we further analyzed the identity of the clipped portion of the sequence following the 3’ end of the exon. To our surprise, most of the clipped sequences were identical to the 5’ end of an earlier exon: e.g., 24.3% of the reads containing the 3’ end of *kl-3* exon 5 were exon-clipped, and of these, 66% were joined to the 5’ end of either exons 2 or 4, suggestive of erroneous back-splicing (Fig 4C, D). Back-splicing is a phenomenon in which the 3’ end of an exon is spliced to the 5’ end of an earlier exon, likely forming a circular RNA molecule (Kristensen, et al. 2019). Such back-spliced products have been identified in a broad range of conditions from healthy tissues to cancer (Conn, et al. 2024). We speculate that such back-splicing may occur if the 3’ end of an exon (at the splice donor site) is unable to locate the correct downstream splice acceptor site (Fig 4C). Similar exon-clipped reads indicative of back-splicing events were observed for *kl-5* exon 19, in which nearly all of the clipped fragments mapped to exons 15 or 17, and *kl-2* exon 6 reads, which mapped to exons 3 and 4 (Fig 4D, Supplementary_file_4).

The presence of exon-clipped reads (instead of experimental artifacts during library preparation) was validated by RT-PCR designed to specifically amplify the correctly spliced products vs. back-spliced products (Fig 5A), which was confirmed by sequencing of the PCR products (Supplementary Fig S3). By further using RT-qPCR with primers to detect correctly spliced vs. back-spliced products, we found that the hybrids had much lower amounts of correctly spliced products (*kl-3* exon 5-exon 6 junction) than the wild type *D. mauritiana*, (Fig 5B, C, Supplementary_file_5). Correspondingly, the hybrids displayed increased levels of back-splicing (the *kl-3* exon 5-exons 4/2 junctions) (Fig 5B, C, Supplementary_file_5), confirming the RNA sequencing results (Fig 4B, D). As an alternative possibility to explain exon-clipped reads, some exons may be duplicated in the genome, leading to the production of RNA reads that appear as back-splicing. Indeed, some exons from the Y-linked gigantic genes are reported to be duplicated (Chang, et al. 2022): however, none but one of ‘back-splicing reads’ detected in our study can be explained by the duplicated exons identified by Chang et al (Chang, et al. 2022). Only one back-splicing event (back-splicing of CCY exon 4 to exon 3) could possibly be explained by a duplicated exon (duplicated exon 3 is located downstream of exon 4) (Supplementary Fig S4). However, analysis of sequence polymorphisms revealed that splicing was between exon 4 and canonical exon 3, demonstrating back-splicing. Thus, these results strongly suggest that ‘exon-clipped’ reads that increase in the hybrids reflect back-splicing events.

**Figure 5.**
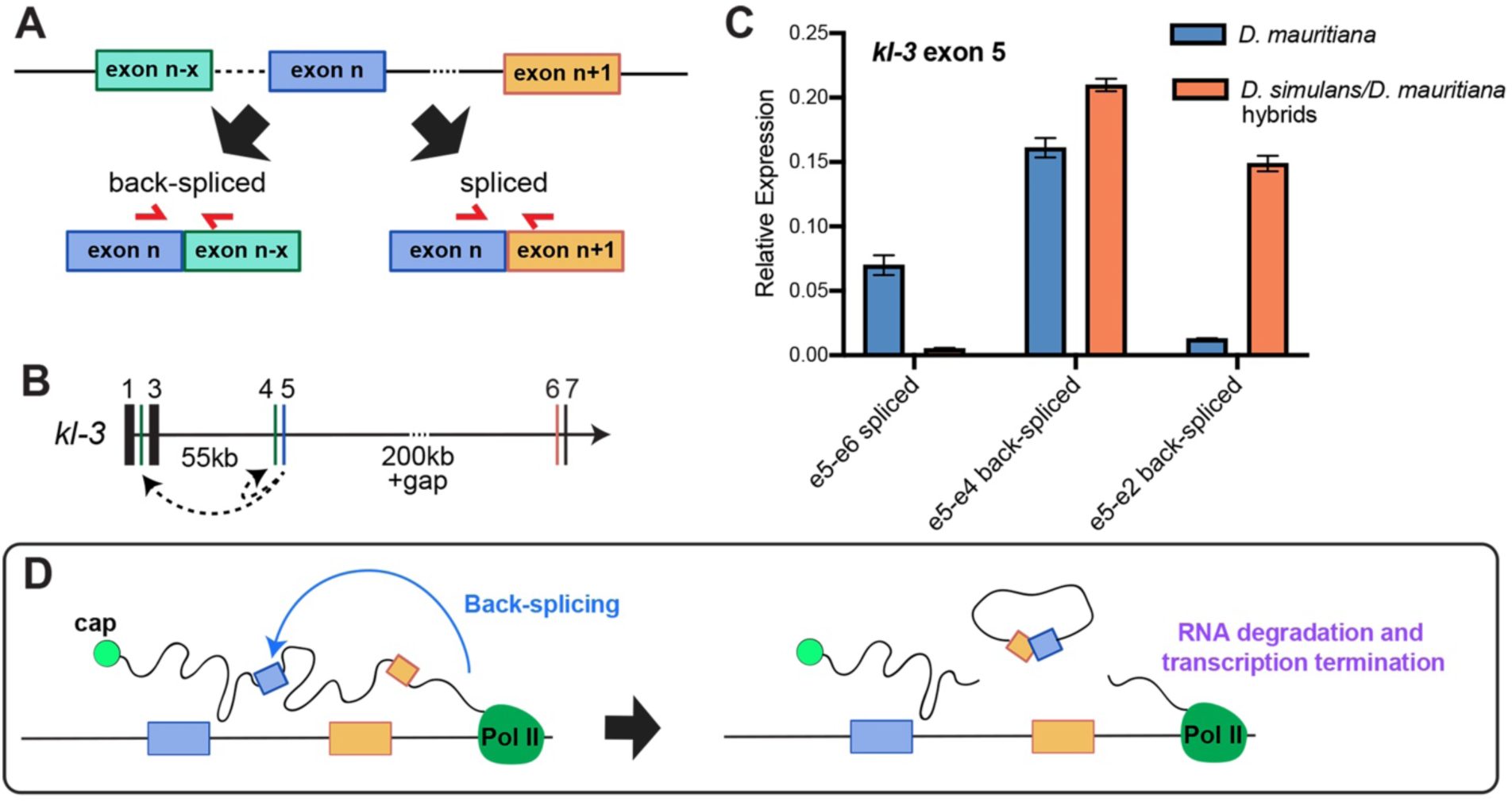
Back-splicing is increased in *D. simulans/D. mauritiana* hybrids. A. RT-qPCR primer designs to detect spliced vs. back-spliced products B. Schematic of the exon-intron structure surrounding kl-3 exon 5 back-splicing junctions. Exons are represented by vertical lines and labeled by exon number. Introns are represented by horizontal lines; a dotted line indicates a gap in the genome assembly C. RT-qPCR to detect the presence of correctly spliced and back-spliced transcripts of kl-3 exon 5, normalized to GAPDH. Error bars represent SEM across three technical replicates. (See Supplementary_file_3 for raw CT values and biological replicates.) D. A model of how back-splicing may lead to transcriptional attenuation (alternative model to the one depicted in Fig. 3b).

It is important to note that exon-clipped reads and back-spliced products were also present in wild type *D. mauritiana* at junctions where the read depth drops, implying that the splicing of gigantic introns is challenging even in the parental species. Importantly, however, mature mRNA was observed in the parental species, suggesting that a sufficient amount of RNAs undergo correct splicing. Back-splicing may also provide an alternative explanation to the drop in RNA sequencing reads observed in the hybrids (Fig 3A): although we have originally speculated that the unspliced gigantic RNA attached to RNA polymerase II may hinder the progression of transcription (Fig 3B)(Fingerhut, et al. 2024), back-splicing will result in uncapped RNA attached to RNA polymerase II, which in turn may cause degradation of nascent RNA and termination of transcription (Fig 5D). Taken together, we conclude that the hybrid males fail to correctly splice gigantic introns, explaining the lack of mature mRNAs of *kl-2*, *kl-3*, and *kl-5* in hybrids (Fig 2). These results indicate that defective splicing of Y-linked fertility genes may contribute to hybrid sterility.

### Splicing defects are a general feature of Y-linked genes with gigantic introns

*kl-2, kl-3,* and *kl-5* are not the only Y-linked genes with gigantic introns. Although other Y-linked gigantic genes did not exhibit clear downregulation in the hybrids (Table 1), several other Y-linked genes displayed drops in read depth within gigantic introns (e.g., *wdy* intron 4, *ppr-Y* intron 1, Fig 6A), suggesting that gigantic introns indeed tend to cause attenuation of transcription. In these cases, the degree of read depth drop was considerably muted compared to *kl-2*, *kl-3*, and *kl-5*, and/or occurred toward the end of the gene (Fig 6A), explaining why the defects were not detected as downregulation of gene expression (Table 1). Nonetheless, in many cases (*WDY, Ppr-y, ORY, CCY, PRY*), gigantic introns were associated with defective splicing in hybrids, encompassing defects similar to *kl-2*, *kl-3*, and *kl-5*, including back-splicing, which was validated by RT-PCR and qPCR (Fig 6B, Supplementary Fig S3F, G, H, Supplementary_file_5). Interestingly, however, defective splicing at the gigantic introns did not necessarily lead to downregulation of transcription (Fig 6A), although defective splicing likely leads to a reduction in functional mRNA. These results imply that splicing failure is a common feature for many gigantic introns.

**Figure 6.**
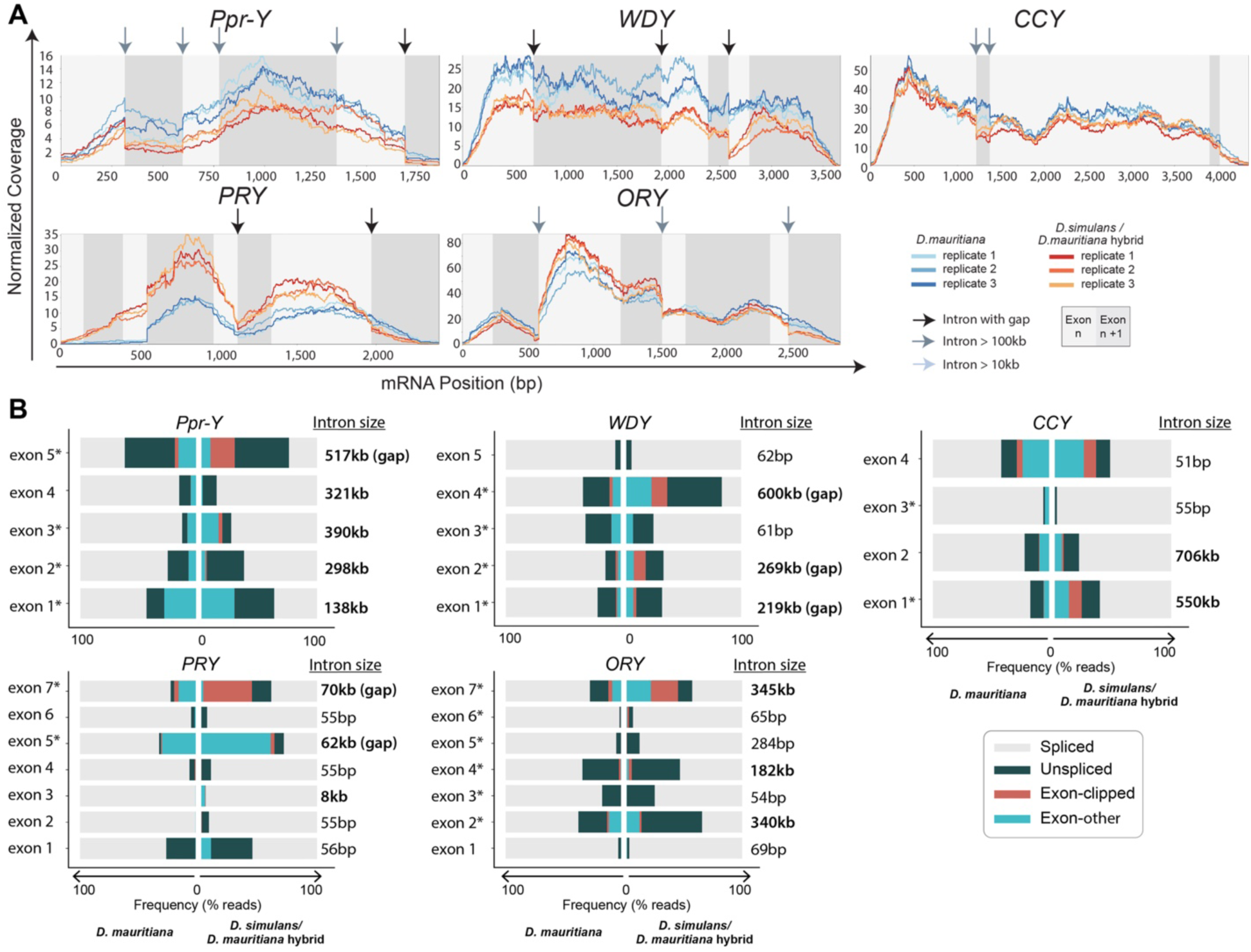
*D. simulans/D. mauritiana* hybrids exhibit splicing defects in Y-linked gigantic genes. A. Normalized read depth coverage of the Y-linked fertility genes (*Ppr-Y, WDY, PRY, ORY, CCY*) in *D. mauritiana* vs. *D. simulans/D. mauritiana* hybrids. B. Sequence read analysis for each exon end of the Y-linked fertility genes (*Ppr-Y, WDY, PRY, ORY, CCY*) in *D. mauritiana* vs. *D. simulans/D. mauritiana* hybrids. Asterisks(*) indicate significant difference in the proportion of correctly spliced reads between *D. mauritiana* and *D. simulans/D. mauritiana* hybrids.

### Size and sequence conservation correlate with splicing errors in hybrids

We found that there is a general correlation between the size of an intron and the degree of its splicing defects (Fig 7, note that for this analysis, we only considered the known length of an intron, ignoring genome assembly gaps). First, we examined the degree of defective splicing for Y-linked genes’ introns, comparing the hybrids to the parental species *D. mauritiana*, which shares the Y chromosomes with the hybrids (Fig 7A). This analysis revealed the trend that larger introns have a higher likelihood of defective splicing, both in the parental species and hybrids [a positive Spearman correlation of 0.50 (p-value 6.3*10^−6^) in the parental species, and 0.55 (p-value 3.33*10^−7^) in the hybrids]. The hybrids exhibited a higher likelihood of defective splicing than the parental species (Fig 7A, Supplementary_file_2). Importantly, the splicing defects in hybrids became more profound as intron size increased, while the parental species showed a lesser increase in such defects. These results imply that although large introns are generally challenging to splice even in parental species, the parental species are able to handle them relatively well. In contrast, the hybrids may be defective in such a mechanism, failing in the splicing of gigantic introns.

**Figure 7.**
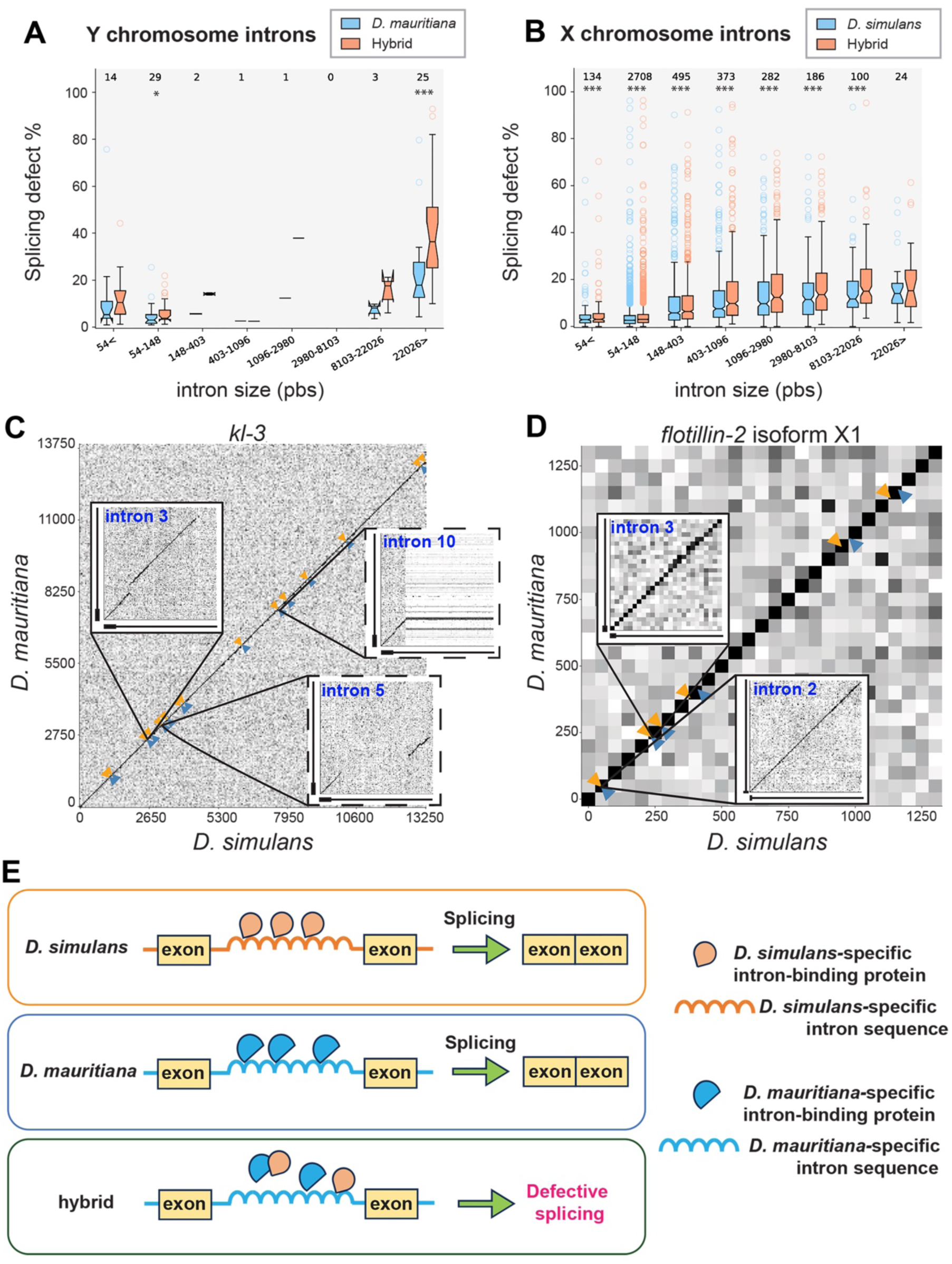
Intron size correlates with splicing defects of Y-linked fertility genes in *D. simulans/D. mauritiana* hybrids. A, B. Frequency of splicing defects (100% - correctly spliced reads/all reads) for Y-linked genes (A) and X-linked genes (B) based on the size of introns. Introns were binned by their size (logarithmic scale with the base of e). Stars indicate statistical significance between *D. simulans/D. mauritiana* hybrids and *D. mauritiana* (Supplementary_file_2) for statistic details.) C, D. Dot plot of sequence conservation between *D. simulans* (X-axis) vs *D. mauritiana* (Y-axis) for *kl-3* gene (Y-linked, C) and *flotillin-2* isoforms X1 gene (X-linked, D). X-axis represent cDNA sequence, with arrows indicating the position of introns. Zoomed-in squares cover the areas containing the end of the preceding exon (up to 500pb, represented by a bold line on the X-Y axes) and the first 5 kb of the intron (represented by a thin line on the X-Y axes). The solid boxes indicate fully assembled introns, and the dotted boxes indicate gap-containing introns. Conservation is indicated by the grey scale: white indicates a conservation (percent of identity) of 50% or less, black indicates a 100% identity. *kl-3* introns 3, 5 and 10 shown in C are 55kb, 270kb and 63kb in *D. mauritiana* and 58kb, 62kb and 21kb in *D. simulans,* respectively. *flotillin-2* introns 2 and 3 in D are 84kb and 1kb for *D. mauritiana* and 90kb and 1kb for *D. simulans*. E. Model of how distinct intronic RNA sequences may cause hybrid sterility. RNA-binding proteins that bind to intronic RNA and aid in the process of splicing are divergent between two species due to distinct intronic sequences. When RNA-binding proteins and intronic RNA sequence are brought together in the context of hybrids, incompatibility between RNA-binding proteins and intronic sequence may cause splicing defects. It is possible that such a factor is X-linked, and such a factor from *D. simulans* cannot correctly splice gigantic genes on *D. mauritiana* Y chromosome. Alternatively, the factor involved in splicing may be autosomal but engage in Dobzhansky-Muller type of dominant negative interactions, interfering with splicing.

We further investigated what aspects of introns influence the splicing defects. First, we asked whether intron size is a determinant of defective splicing for genes linked to the X chromosome and autosomes. When we analyzed splicing defects on X-linked genes, comparing the hybrids and *D. simulans*, the hybrids exhibited more splicing defects than *D. simulans*, implying widespread splicing defects in the hybrids (Fig. 7B, Supplementary_file_2). However, the relative difference in splicing defects between *D. simulans* and the hybrid was fairly constant across intron sizes (Fig. 7B, Supplementary_file_2). Autosomal genes also exhibited the same trend as X-linked genes: the hybrid exhibited splicing defects compared to *D. simulans*, but the defects did not become more pronounced with increasing intron sizes (Supplementary Figure S5), as observed with the Y-linked genes (Fig 7A, Supplementary_file_2). However, given that Y-linked genes are much larger than X-linked or autosomal genes (the largest intron from X-linked/autosomal genes is 106kb from the gene, *rg*, compared to >∼400kb plus a gap for the Y-linked genes), it may simply suggest that intron size is a major parameter of defective splicing in the hybrids.

However, further analysis indicated that introns of Y-linked genes may also have other features, in addition to size, that make them sensitive to splicing defects. The analysis of DNA sequence homology of the *kl-3* gene revealed that its introns are highly divergent between *D. simulans* and *D. mauritiana*, whereas the exon sequences are well conserved (Fig. 7C). Other gigantic introns of the Y-linked genes are also poorly conserved, whereas small introns of the Y-linked genes are generally well conserved (Supplementary Figure S6, Supplementary_file_6). In contrast, both exons and introns of *Flotillin-2*, an example of an X-linked large gene, are highly conserved between two species (Fig. 7D). The largest introns from the X-linked and autosomal genes (there are only three genes with large introns that are expressed in the testis) are well conserved (Supplementary Figure S7), and the hybrids did not exhibit enhanced splicing defects at these introns (Supplementary Figure S8). The genome-wide analysis of sequence conservation further revealed that the introns of Y-linked genes exhibit marked divergence between *D. simulans* and *D. mauritiana,* whereas exons of Y-linked genes, as well as both introns and exons of the X chromosome/autosomes, are highly conserved (Supplementary Figure S6, Supplementary_file_6, Methods). In conclusion, gigantic introns of the Y-linked genes are far larger than the largest introns of X-linked and autosomal genes. Moreover, these gigantic introns of the Y-linked genes contain gaps and are not conserved, whereas introns of X-linked and autosomal genes are well conserved. It is important to note that small introns of the Y-linked genes are well conserved, and do not exhibit splicing defects (Fig. 4, Fig. 7A, Supplementary_file_2). Together, we propose that the size and/or poor conservation of the introns contribute to the splicing defects of Y-linked genes in the hybrids. Further investigations are required to establish the causal link between intron’s characteristics (e.g. size, conservation) and failure in splicing in hybrids.

## Discussion

In this study, we show that the splicing of Y-linked gigantic genes is severely impacted in male hybrids between *D. simulans* and *D. mauritiana*, leading to the downregulation of multiple fertility genes. We propose that such downregulation of fertility genes contributes to hybrid sterility, together with other proposed causes of hybrid male sterility. Our work provides a potential explanation as to why the heterospecific Y chromosome does not support male fertility. The Y chromosome is the most rapidly diverging chromosome across species (Hughes, et al. 2012; Soh, et al. 2014; Hughes and Page 2015; Kotov, et al. 2022), thus our finding may provide a generalizable principle of Y chromosome-mediated hybrid sterility.

Previous studies have provided significant insights into the mechanisms of hybrid male sterility, and the presence of multiple hybrid incompatibility genes was shown (see Introduction). More broadly, one of the leading explanations for hybrid sterility is that genomic conflicts caused by independently evolved male meiotic drivers result in hybrid sterility (Frank 1991; Hurst and Pomiankowski 1991; McDermott and Noor 2010). Indeed, there are several examples where male sterility and meiotic drive are caused by the same genetic elements, including *tmy* in *D. simulans/D. mauritiana* (Tao, et al. 2001), *Overdrive* in *D. pseudoobscura* (Phadnis and Orr 2009), and *Sex Ratio (SR)* in *D. pseudoobscura* (Bladen, et al. 2024). Another proposed cause of hybrid sterility is transposon derepression (Kelleher, et al. 2012; Dion-Cote, et al. 2014; Parhad, et al. 2017; Castillo and Moyle 2022; Kotov, et al. 2024), although there are cases where transposons are not dramatically derepressed in sterile hybrids (e.g., hybrids between *D. arizonae* and *D. mojavensis*) (Banho, et al. 2021). Our RNA sequencing analysis revealed some degrees of transposon derepression in the *D. simulans/D. mauritiana* hybrid males (Supplementary Table S1), which may also contribute to hybrid dysfunction.

The present study demonstrates the defective splicing of gigantic introns of Y-linked fertility genes in *D. simulans/D. mauritiana* hybrids, potentially providing a link between rapid divergence of the Y chromosome and hybrid male sterility. Previous studies have shown that gigantic introns of Y-linked fertility genes pose challenges in the processes of gene expression, such as transcription and/or splicing, even in wild type flies, requiring additional factors to aid in the process: multiple proteins have been identified to bind to intronic transcripts (e.g., Blanks, Heph, and Maca) and are required for the production of mature mRNA from fertility genes with gigantic introns (Fingerhut, et al. 2019; Zhu and Fukunaga 2021). Additionally, the transcription of gigantic genes is sensitive to the perturbation of splicing (Fingerhut, et al. 2024), further illuminating the challenging nature of producing mRNA from gigantic genes. Because sequences of the Y-linked gigantic introns are highly divergent (Supplementary Figure S6), it is conceivable that RNA-binding proteins, which bind gigantic introns to facilitate their splicing, become incompatible between species (Fig 7E). Indeed, three proteins that have been shown to bind to satellite RNA from gigantic introns - Blanks, Heph and Maca (Fingerhut, et al. 2019; Zhu and Fukunaga 2021) - exhibit a high degree of sequence divergence (Supplementary Table S1) More specifically, gigantic introns are likely spliced recursively, which may involve proteins that bind intronic sequence. Because intronic sequences are highly divergent between *D. simulans* and *D. mauritiana* (Fig 7C), intron binding proteins may become incompatible. It is tempting to speculate that incompatibility of this nature contributes to Haldane’s rule (Haldane 1922), specifically impacting male fertility.

In summary, the present study provides a new model on a cause of hybrid male sterility, providing a potential molecular explanation as to why the Y chromosomes are incompatible between species.

## Methods

### Fly husbandry and strains used

All *Drosophila melanogaster* strains were raised on standard Bloomington medium at 25 °C. The following stocks were used: *D. simulans* wXD1 and *D. mauritiana* w12 (obtained from Dr. Ching-Ho Chang).

### Immunofluorescence staining

Testes from 1- to 3-day-old males were dissected in 1x PBS and fixed in 4% formaldehyde in 1x PBS for 30 minutes. Fixed testes were then washed in 1x PBST (PBS containing 0.1% Triton X-100) for at least 1.5 hours, followed by incubation with primary antibodies diluted in 1x PBST containing 3% BSA at 4 °C overnight. Samples were washed three times in 1x PBST for 30 minutes each and then incubated with secondary antibodies in 1x PBST with 3% BSA at 4 °C overnight. After a similar washing procedure, samples were mounted in VECTASHIELD with DAPI (Vector Labs). Images were acquired using a Leica Stellaris 8 confocal microscope with a 63x oil immersion objective lens (numerical aperture 1.4) and processed with Fiji (ImageJ) software. The primary antibodies used were: anti-Pontin (1:200; guinea pig) (Fingerhut and Yamashita 2020), anti-pH3 ser10 (1:200, rabbit), and anti-ATP5a (1:1000; mouse; Abcam, ab14748). Phalloidin-Alexa Fluor 488 (1:200; Thermo Fisher Scientific, A12379) was used to stain F-actin. Alexa Fluor-conjugated secondary antibodies (Life Technologies) were used at a 1:200 dilution.

### Single-molecule RNA Fluorescent *in situ* hybridization

RNA FISH was performed as previously described (Fingerhut and Yamashita 2023). All solutions used were RNase free. Testes from 1–3 day old flies were dissected in 1X PBS and fixed in 4% formaldehyde in 1X PBS for 30 minutes. Testes were washed briefly in 1X PBS and permeabilized in 70% ethanol overnight at 4°C. Testes were briefly rinsed with wash buffer (2X saline-sodium citrate (SSC), 10% formamide) and then hybridized overnight at 37°C in hybridization buffer (2X SSC, 10% dextran sulfate (sigma, D8906), 1mg/mL E. coli tRNA (sigma, R8759), 2mM Vanadyl Ribonucleoside complex (NEB S142), 0.5% BSA (Ambion, AM2618), 10% formamide). Following hybridization, samples were washed three times in wash buffer for 20 minutes each at 37°C and mounted in VECTASHIELD with DAPI (Vector Labs). Images were acquired using a Leica Stellaris8 confocal microscope with a 63X oil immersion objective lens (NA = 1.4) and processed using Fiji (ImageJ) software.

Fluorescently labeled probes were added to the hybridization buffer to a final concentration 100nM. Probes against *kl-3*, *kl-5*, and *kl-2* exons were designed using the Stellaris RNA FISH Probe Designer (Biosearch Technologies, Inc.) available online at www.biosearchtech.com/stellarisdesigner. Each set of custom Stellaris RNA FISH probes was labeled with Quasar 670, Quasar 570 or Fluorescein-C3. Probe information can be found in Supplementary_file_7 (Fingerhut, et al. 2019).

### RNA isolation and sequencing

Total RNA was purified from 2-5 day old adult testes (100 pairs/sample) by TRIzol (Invitrogen) extraction according to the manufacturer’s instructions. Libraries were prepared for RNA sequencing using the KAPA Biosystems RNA HyperPrep Kit with RiboErase according to manufacturer’s directions with some modifications. Briefly, 500 ng of total RNA was ribo-depleted by hybridization of complementary DNA oligonucleotides. The set of complementary oligonucleotides was a custom panel designed for *Drosophila*. This was followed by treatment with RNase H and DNase to remove rRNA duplexed to DNA and original DNA oligonucleotides. The enriched fraction was then fragmented with heat and magnesium, and first-strand cDNA was generated using random primers. Strand specificity was achieved during second-strand cDNA synthesis by replacing dTTP with dUTP, which quenches the second strand during amplification, and the cDNA is then A-Tailed. The final double strand cDNA was then ligated with indexed adapters. Finally, the library was amplified using a DNA Polymerase that cannot incorporate past dUTPs, effectively quenching the second strand during PCR. Libraries were enriched for fragments between 500 – 1000bp with two additional cycles of PCR followed by a size selection using a 1.5% gel on a Pippin Prep (Sage Science) electrophoresis instrument. Final libraries were quantified by qPCR and Fragment Analyzer. Samples were sequenced on a NOVASEQ 6000, producing 250 × 250bp paired-end reads.

### RT-PCR, RT-qPCR, and sequencing

Total RNA was purified from 2-5 day old adult testes (50 pairs/sample) by TRIzol (Invitrogen) extraction according to the manufacturer’s instructions. 1 µg total RNA was reverse transcribed with SuperScript III First-Strand Synthesis Supermix for qRT-PCR (Invitrogen), followed by PCR using Phusion High Fidelity DNA Polymerase (New England Biolabs) with DMSO according to the manufacturer’s instructions. PCR products were purified via gel extraction using a QIAquick Gel Extraction Kit (Qiagen). All reactions were done in technical duplicates with two biological replicates. Exon junctions were chosen based on the most highly abundant back-spliced transcripts from our RNAseq data set. PCR primers were designed to span exon junctions such that the product would only be detected if the 3’ end of the first exon was spliced directly to the 5’ end of the second exon. Primer sequences are listed in Supplementary_file_7. DNA concentration was measured by a Qubit 3.0 Fluorometer (Invitrogen). One replicate of *D. mauritiana* and one replicate of *D. simulans/D. mauritiana* hybrid PCR products were sequenced for each PCR target. Premium PCR Sequencing was performed by Plasmidsaurus using Oxford Nanopore Technology with custom analysis and annotation.

For RT-qPCR, total RNA was isolated and reverse transcribed as described above. qPCR was performed using SYBR Green PCR Master Mix (Applied Biosystems) on a QuantStudio 6 Flex Real-Time PCR system (Applied Biosystems). Relative expression levels were normalized to GAPDH and *D. mauritiana* controls. All reactions were done in technical triplicates with two biological replicates. Graphical representation is inclusive of all replicates. Primer sequences are listed in Supplementary_file_7.

### Bioinformatics analysis

#### Read alignment

Paired-end reads (251 x 251 bp) were quality trimmed using fastp with the following options: “--length_required 100 --disable_adapter_trimming --trim_poly_g --cut_tail -M 20”. We used the *D. simulans* refseq genome (GCF_016746395.2) and the long read assembly of *D. mauritiana* (https://doi.org/10.5061/dryad.280gb5mr6). Alignment was performed using STAR (2.7.10a_alpha_220818) with “--alignIntronMa× 1000000 --twopassMode Basic” parameters.

#### Y gene annotation

MMseqs2 (version 2fad714b525f1975b62c2d2b5aff28274ad57466) was used to align the translated products of *D. melanogaster* Y genes to both the *D. mauritiana* and *D. simulans* genomes (https://doi.org/10.5061/dryad.280gb5mr6), with the parameters “-s 7.5 --max-seqs 1000 -a”. Similar to Chang et al.(Chang, et al. 2022), we manually parsed the tabulated alignment file to annotate putative exon positions. Our RNA-seq data were considerably deeper than the data set that was utilized by Chang et al, allowing us to further refine the annotation (our sequencing data contained 129, 64, or 71 billion nucleotides (after quality trimming) for hybrid, *D. simulans* and *D. mauritiana*, respectively. Chang et al. used the sequencing data from (Lin, et al. 2018) and (Chakraborty, et al. 2021), which contained 15 and 14 billion nucleotides (after quality trimming) for *D. simulans* and *D. mauritiana*, respectively). This resulted in the identification of previously unannotated exons: we found a couple of exons in *D. melanogaster* that have been split into multiple exons in *D. mauritiana*. We corrected the exon boundary, such that it matched the observed read junctions. Finally, we resolved exon duplications by ensuring that all predicted exons have expected junction reads (except for *kl-3* exon 1, which we couldn’t resolve) (Supplementary_file_1). Lastly, we validated our results by ensuring that the transcripts we annotated encode in-frame peptides.

#### Transposable element

RepeatMasker (http://www.repeatmasker.org) was used to extract the transposable elements using One Code To find Them All from both *D. simulans* and *D. mauritiana* (https://doi.org/10.5061/dryad.280gb5mr6). We merged the sequences of the transposable elements from both species. Then we used TEtools (Lerat, et al. 2017) to count the reads matching to transposable elements. Finally, we used DESeq2 (Love, et al. 2014) to test for differential expression of transposable elements.

#### Splicing defect

We Used OmniSplice (Lannes, et al. 2025) (https://github.com/rLannes/OmniSplice/releases/tag/Revision) to identify splicing defects. For the statistical analyses, we used the “statsmodels” library in Python. To identify junctions that are differently spliced between *D. mauritiana* and *D. simulans/D. mauritiana* hybrids (Figure 4 and 6 bar-plot), we used a linear binomial model “smf.glm (’successes + failures ∼ group’, family=sm.families.Binomial(), data=df).fit()” with “successes” being the Spliced category and “failures” being the sum of the “Unspliced”, Clipped” and “Exon_Other” categories. For Figure 7 bar-plots, we used the Wilcoxon signed rank test to determine whether splicing efficiency is different within each intron size group between conditions. We corrected p-values with the Benjamini/Hochberg method using the “fdrcorrection” function. All calculated p-values and corrected p-values are included in Supplementary_file_2

#### Genome wide intron and exon sequence identity estimates

Because the sequences of gigantic introns were poorly conserved between *D. simulans* and *D. mauritiana*, it was not possible to calculate the homology of intronic sequences. Thus, using bowtie2, DNA sequencing of *D. simulans* (SRR22548176) was aligned to both the genomes of *D. simulans* and *D. mauritiana* (long reads genome assembly (https://doi.org/10.5061/dryad.280gb5mr6)). The proportion of bases covered by at least one read was used as a proxy of sequence conservation (because poorly conserved sequences will not be mapped). Comparison between *D. simulans* short read sequencing vs. *D. simulans* long-read assembled genome served as the control, where all sequence reads must be perfectly conserved (however, note that some sequences showed low coverage, likely because of genome assembly/strain difference). For the X-linked genes, we used the gtf file of the *D. simulans* genome assembly (https://doi.org/10.5061/dryad.280gb5mr6) and for the Y-linked genes, we used our annotation. Then, using those gtf files, we made two bed files: one describing all exons and the other all introns. Finally, using bedtools coverage (version v2.29.2)(Quinlan and Hall 2010), we computed the percentage of bases covered by at least one read for both all introns and all exons. (see GitHub repository to reproduce the code: https://github.com/rLannes/Fontan_2025)

#### Differential expression

We performed two analyses. For the Y genes, we aligned both *D. simulans* and *D. simulans/D. mauritiana* hybrids to the *D. mauritiana* refseq genome. And for the X and autosome genes, we aligned both *D. simulans* and *D. simulans/D. mauritiana* hybrids to the *D. simulans* refseq genome. Then, using feature counts (-s 2 -p -B options) and Deseq2, we performed differential gene expression analyses. Analysis of the RNA sequencing results confirmed downregulation of several autosomal genes in hybrids, consistent with previous studies (Michalak and Noor 2003, 2004; Moehring, et al. 2007; Catron and Noor 2008; Sundararajan and Civetta 2011): *dj, Mst98Ca*, and *Mst84Dc* (which is annotated as a lncRNA LOC27208299 in the recent *D. simulans* genome annotation (GCF_016746395.2)) (estimated log2 fold change −1.37, −0.81, −1.03, and adjusted p-value 7.8*10^−51^, 1.54*10^−14^, 0.07 for *dj*, *Mst98Ca*, and *LOC27208299,* respectively).

#### Plotting

All figures were generated using python (3.8.10) and matplotlib (3.7.1). The dot plot was generated using a custom code available on GitHub. Briefly, we first cut all sequences into smaller chunks. Using the Needleman-Wunsch algorithm we aligned all the chunks of a sequence to all the chunks of the other sequence. We used the ECDNA matrix for scoring with the following parameters (gap opening: −10, gap extension: −0.5). From those alignments, we reconstructed the dot plot. For the coverage plots, we designed an in-house code available at https://github.com/rLannes/Fontan_2025.

## Data availability

All data are provided in the manuscript. RNA sequencing data is deposited to SRA under the bioproject accession number: PRJNA1248387.

## Code Availability

Bioinformatics code used in this study is available at https://github.com/rLannes/Fontan_2025.

## Acknowledgements

We thank the members of the Yamashita lab for discussions and comments on the manuscript, and Phil Sharp for helpful suggestions. We thank Flybase and the National Drosophila Species Stock Center for reagents and critical information. We thank the Genome Technology Core at the Whitehead Institute for their consultation and aid in designing and performing RNA sequencing experiments.

## Author contributions

Conceptualization: AF, RL, YY

Methodology: AF, RL, JFi, JFl, YY

Investigation: AF, RL, JFi, JFl, YY

Funding acquisition: YY

Supervision: YY

Writing and editing: AF, RL, JFi, YY

