## Supplementary Figures for "Defective splicing of Y-chromosome-linked gigantic genes contributes to hybrid male sterility in Drosophila"

### Supplementary Figures and Table

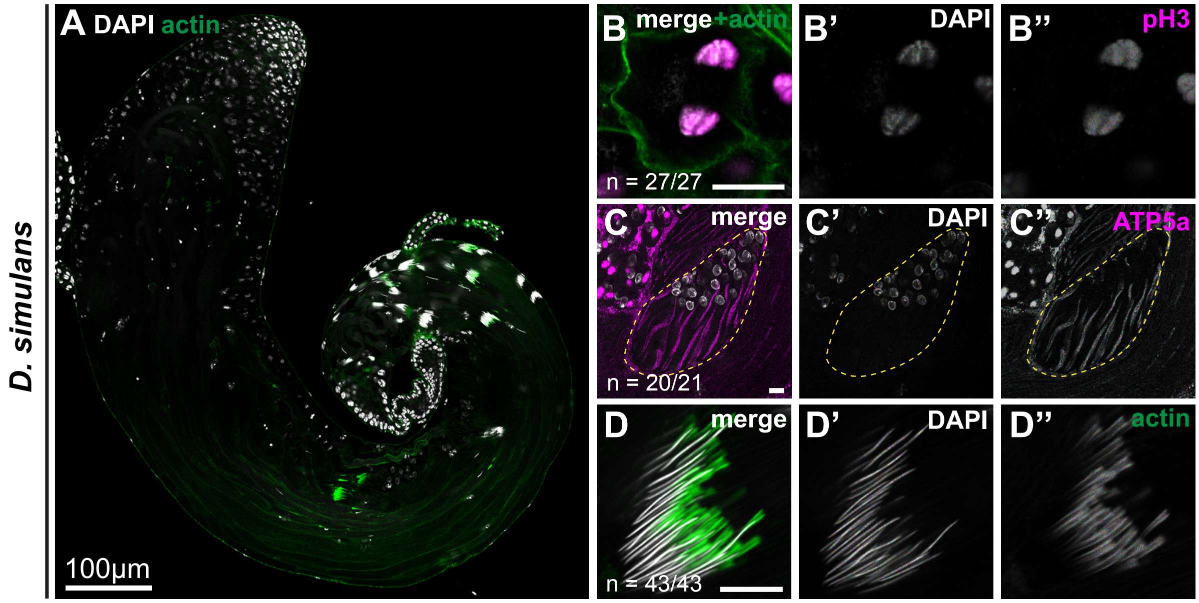


**Supplementary Figure S1: *D. simulans* spermatogenesis**

A-D. *D. simulans* testis and germ cells. Low magnification view of the whole testis (A) stained for F-actin (green, stained with phalloidin-Alexa 488) and DAPI (gray). Meiotic anaphase (B) stained for phospho-H3 ser10 (magenta) and DAPI (gray). Early elongating spermatids (C) stained for ATP5a (magenta, mitochondria) and DAPI (gray). Needle stage spermatids (D) stained for F-actin (green, stained with phalloidin-Alexa 488, individualization complex). Scale bar: 10µm, unless otherwise noted.

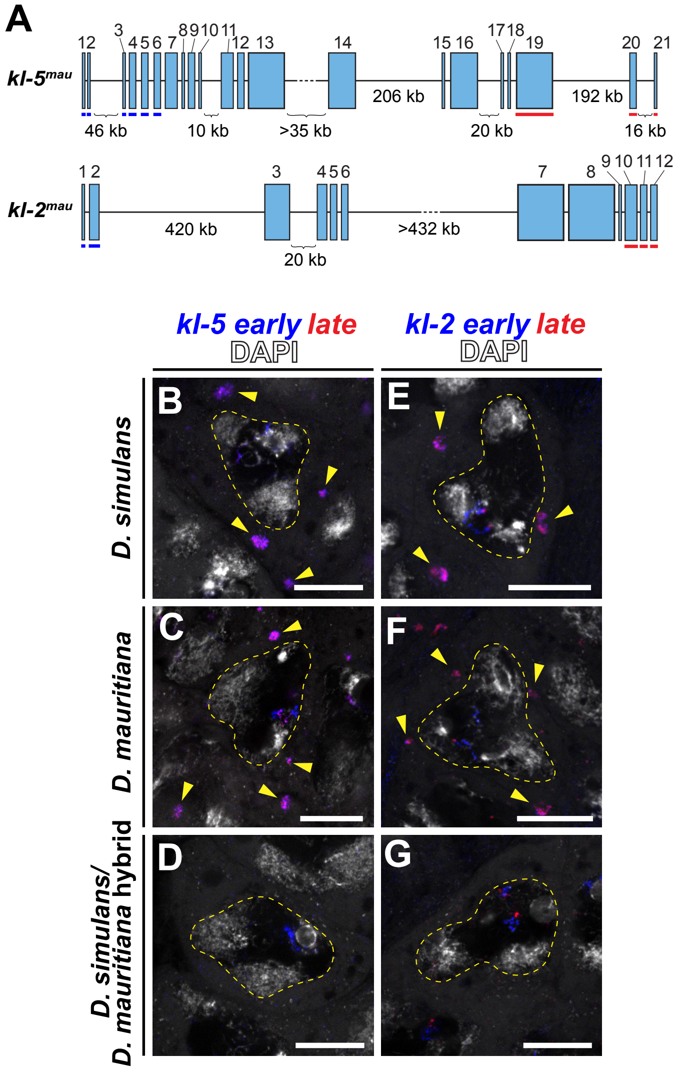


**Supplementary Figure S2: Expression of *kl-2* and *kl-5* in the spermatocytes of the hybrids and parental species.**

A. *D. mauritiana kl-5* (*kl-5$^{mau}$*) and *kl-2* (*kl-2$^{mau}$*) gene structure. The positions of smRNA FISH probes are indicated.

B-F. RNA in situ hybridization for *kl-5* transcripts in late-stage spermatocyte nuclei of *D. simulans* (B), *D. mauritiana* (C) and *D. simulans/D. mauritiana* hybrids (D). Spermatocyte nuclei are indicated by dashed lines. Arrowheads indicate cytoplasmic kl-granule containing mature *kl-5* mRNA. Blue: *kl-5* early exons. Red: *kl-5* late exons. Gray: DAPI. Scale bar: 10µm.

E-G. RNA in situ hybridization for *kl-2* transcripts in late-stage spermatocyte nuclei of *D. simulans* (E), *D. mauritiana* (F) and *D. simulans/D. mauritiana* hybrids (G). Spermatocyte nuclei are indicated by dashed lines. Arrowheads indicate cytoplasmic kl-granule containing mature *kl-2* mRNA. Blue: *kl-2* early exons. Red: *kl-2* late exons. Gray: DAPI. Scale bar: 10µm.

A

| RT-PCR Target | Amplicon size (bp) |
|---|---|
| *kl-3* exon 5-exon 6 | 512 |
| *kl-3* exon 5-exon 4 | 506 |
| *kl-3* exon 5-exon 2 | 410 |
| *kl-5* exon 19-exon 20 | 554 |
| *kl-5* exon 19-exon 17 | 504 |
| *ory* exon 7-exon 8 | 398 |
| *ory* exon 7-exon 5 | 301 |
| *ccy* exon 4-exon 5 | 422 |
| *ccy* exon 4-exon 3 | 600 |

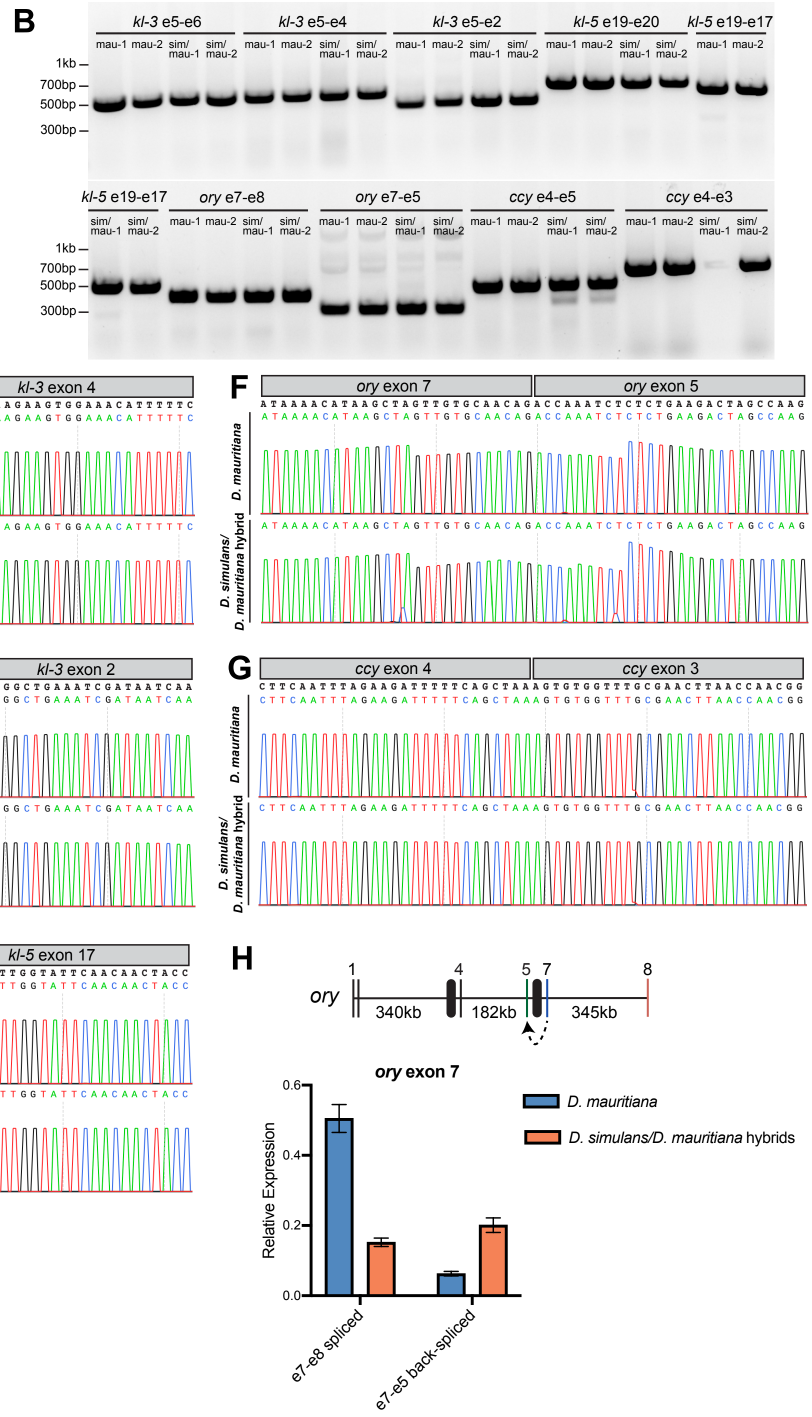


**Supplementary Figure S3: Validation of back-splicing products.**

A-B. RT-PCR for back-spliced transcripts. (A) List of expected PCR amplicon sizes. Primer sequences can be found in Supplementary_file_3. (B) RT-PCR products for two biological replicates of each *D. mauritiana* and *D. simulans/D. mauritiana* hybrids. *CCY* e4-e3 failed to amplify in one of the hybrid replicates.

C-G. Oxford Nanopore sequencing of the PCR products generated in (A-B) aligned to the corresponding exon-exon junction. Only the sequence immediately around the junction is shown.

H. RT-qPCR results of spliced vs. back-spliced products involving *ORY* exon 7. Schematic of the exon-intron structure surrounding *ory* exon 7 is shown in the upper panel. Exons are represented by vertical lines and labeled by exon number. Introns are represented by horizontal lines; a dotted line indicates a gap in the genome assembly. RT-qPCR results were normalized to GAPDH. Error bars represent SEM across three technical replicates. (See Supplementary_file_3 for raw CT values and biological replicates.)

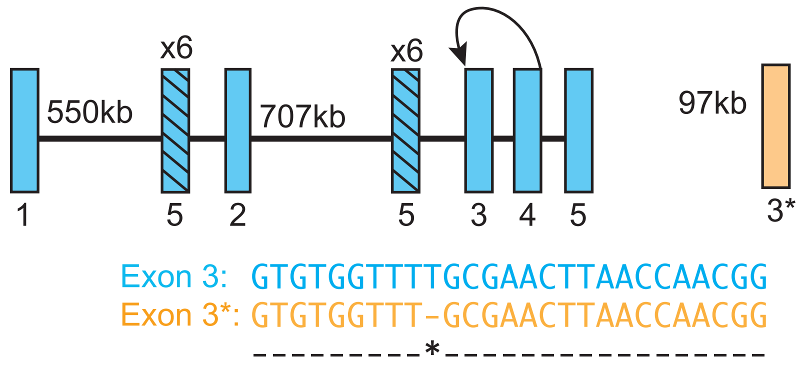


**Supplementary Figure S4: Structure of CCY and its back-splicing patterns**
CCY gene contains multiple duplicated exons. Exon 4 - exon 3 spliced product was detected, and this may be explained by exon 4 - exon 3 back-splicing or exon 4- exon 3* forward-splicing (exon 3* is located 97 kb downstream of CCY's terminal stop codon). However, based on SNP (shown at the bottom), it was determined that exon 4 – exon 3 back-splicing

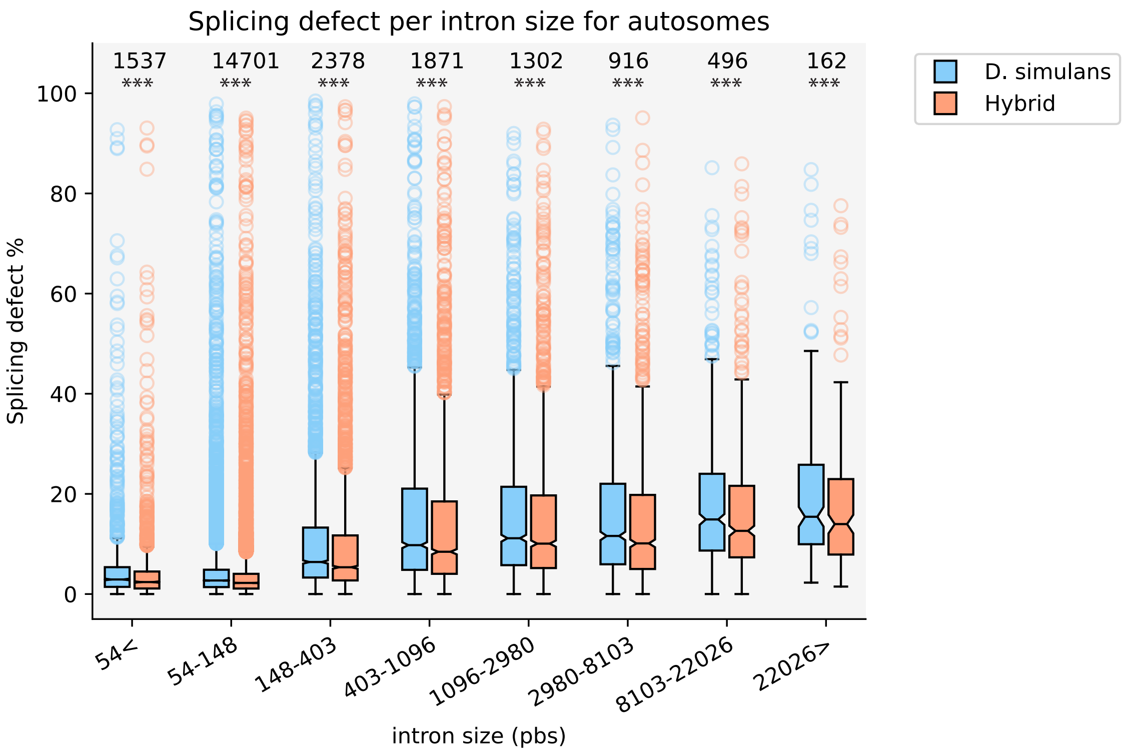


**Supplementary Figure S5: Frequency of splicing defects for autosomal genes.**
Frequency of splicing defects (100% - correctly spliced reads/all reads) for autosomal genes based on the size of introns. Introns were binned by their size (logarithmic scale with the base of e). Stars indicate statistical significance between *D. simulans/D. mauritiana* hybrids and *D. mauritiana* (Supplementary_file_2 for statistics details.)

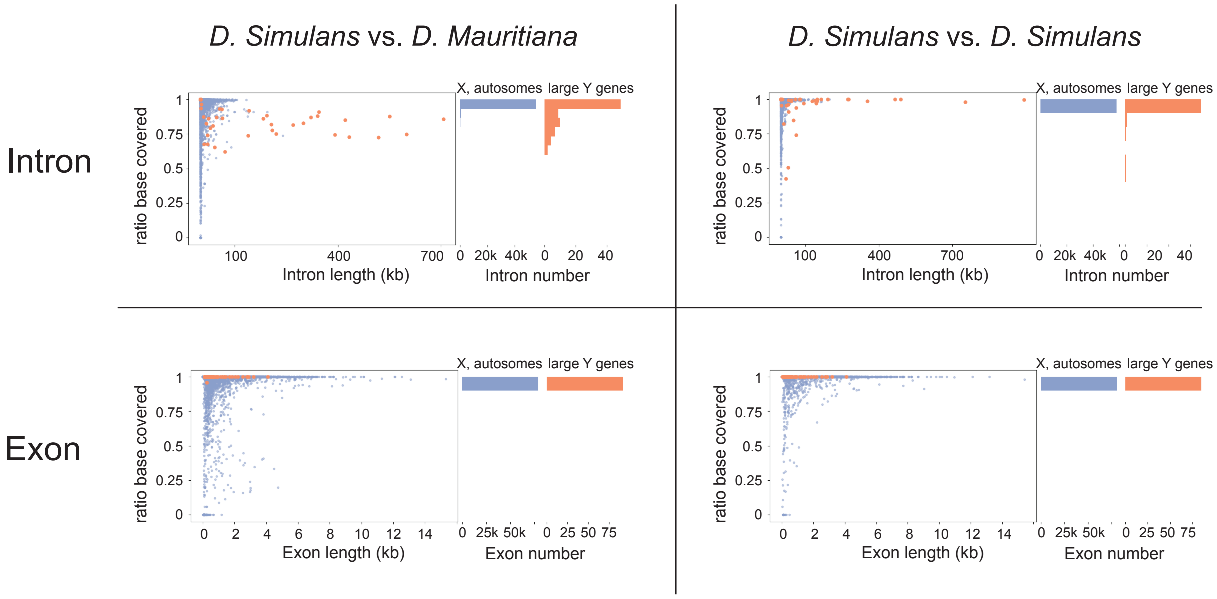


**Supplementary Figure S6: estimation of DNA sequence conservation between *D. simulans* and *D. mauritiana* for exons and introns**

Ratio of covered bases for genomic features (intron: top row, exon: bottom row) by short read DNA sequencing from *D. simulans* aligned against either *D. mauritiana* genome (left) or *D. simulans* genome (right). Features from X and autosomes are shown in blue, and large Y genes in orange. Histograms (right to each scatter plot) indicate the distribution of introns based on the conservation (Y axis).

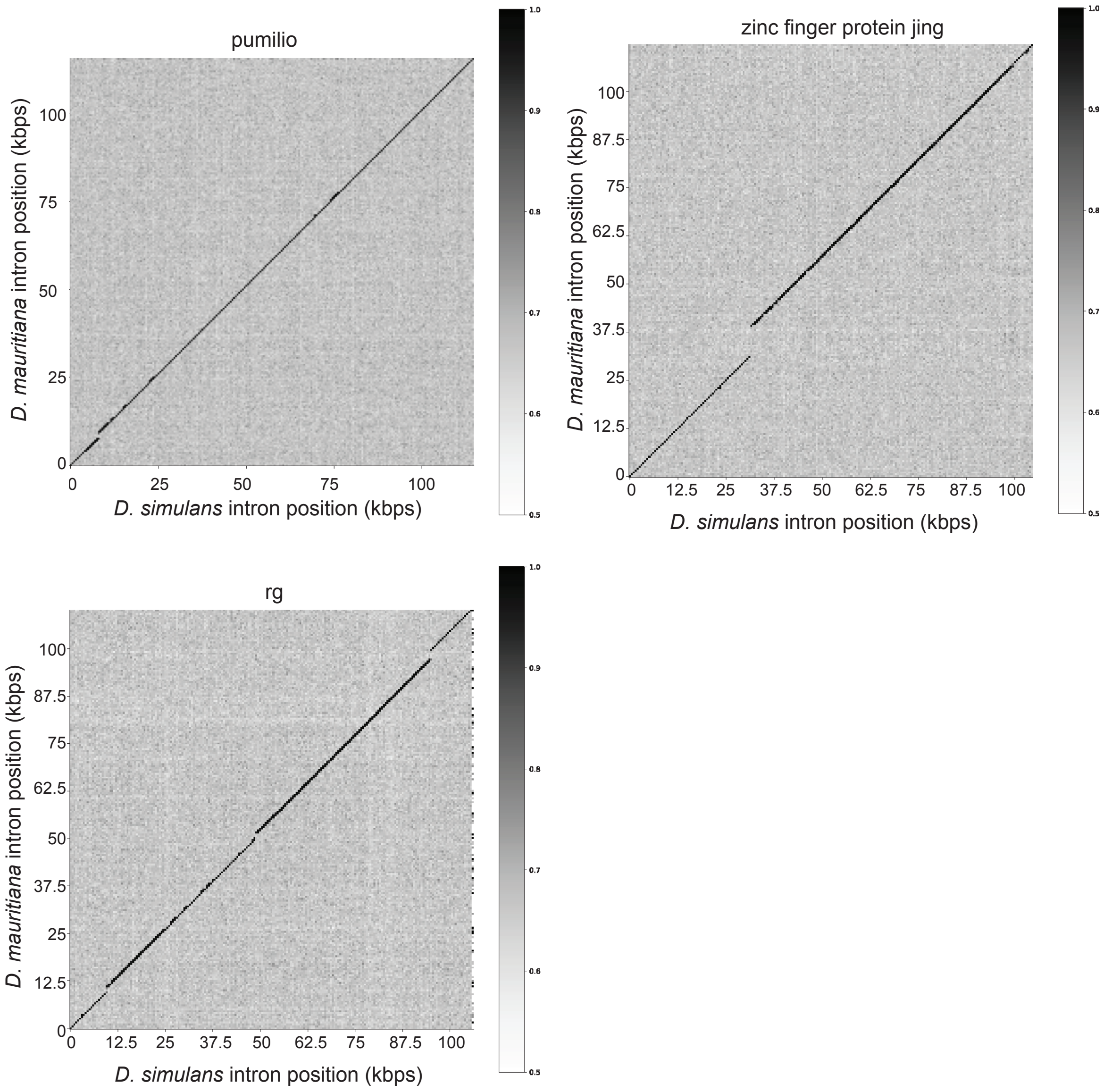


**Supplementary Figure S7: Large introns of autosomal genes are conserved.**
There are only three autosomal genes that have large introns (>100kbps) and are expressed in the testes. They are well-conserved. Window size is 500 bps, and alignment was performed with the Needleman-Wunsch algorithm. Scoring: ECDNA matrix, gap open -10 and gal extension -0.5.

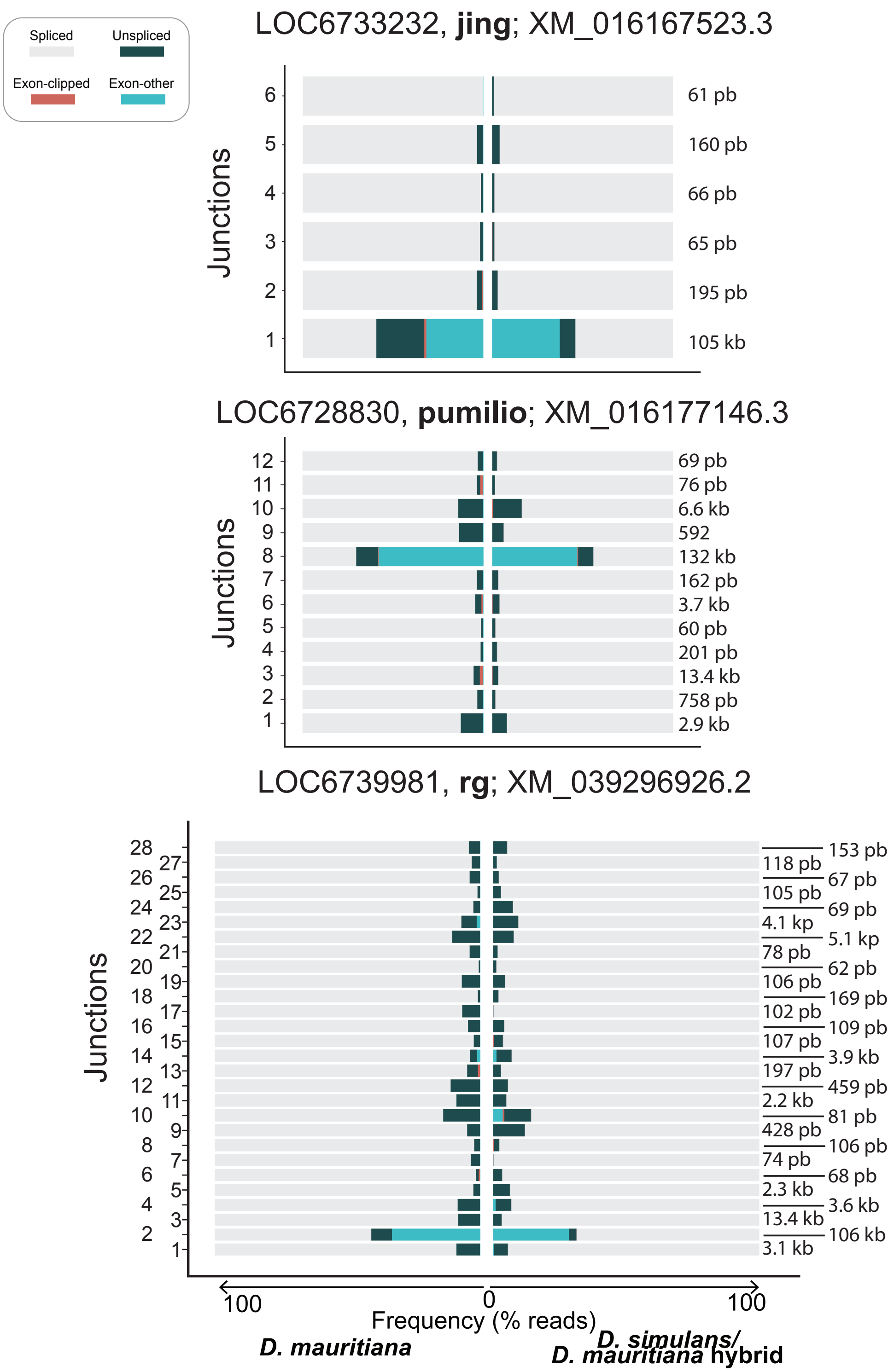


**Supplementary Figure S8:** Sequence read analysis for each exon end of the three genes with large introns linked to the X chromosome or autosomes. Large introns of these genes did not exhibit a statistically significant difference between *D. mauritiana* and *D. simulans/D. mauritiana* hybrids.

**Blanks**

| | Dmel | Dsim | Dmau | Dsec |
|---|---|---|---|---|
| Dmel | - | 54.87 | 54.97 | 57.34 |
| Dsim | - | - | 91.94 | 54.73 |
| Dmau | - | - | - | 53.39 |
| Dsec | - | - | - | - |

**Heph**

| | Dmel | Dsim | Dmau | Dsec |
|---|---|---|---|---|
| Dmel | - | 66.52 | 73.39 | 94.44 |
| Dsim | - | - | 89 | 67.18 |
| Dmau | - | - | - | 74 |
| Dsec | - | - | - | - |

**Maca**

| | Dmel | Dsim | Dmau | Dsec |
|---|---|---|---|---|
| Dmel | - | 78.09 | 79.68 | 78.88 |
| Dsim | - | - | 94.72 | 95.08 |
| Dmau | - | - | - | 96.34 |
| Dsec | - | - | - | - |

**Supplementary Table S1: Protein sequence divergence of Blanks, Heph and Maca**
% identity from amino acid alignment using blosum62 and gap open -10 and gap extension 0.5
